# PZLS-1/PDZRN controls muscle polarity downstream of Wnt-Ror-Dvl signalling in *C. elegans* muscles

**DOI:** 10.64898/2026.10.09.757662

**Authors:** Elise Cheynet, Sandra Duperrier, Thomas Boulin

## Abstract

Non-canonical Wnt pathways provide directional cues for cell polarity, but the mechanisms that translate these cues into asymmetric cellular organisation remain incompletely defined. We previously showed that a non-canonical EGL-20/Wnt-CAM-1/Ror-DSH-1/Dishevelled pathway establishes polarised membrane compartments in *Caenorhabditis elegans* body wall muscles. However, how EGL-20/Wnt signalling is translated into muscle polarity remains unknown. Here, we identify PZLS-1, the *C. elegans* ortholog of human PDZRN3, PDZRN4 and PDZD4, as a regulator of muscle polarity. Although PDZRN proteins are known targets of non-canonical Wnt-Ror signalling, how they are spatially organised within polarised cells and how their protein domains contribute to their function remain largely unknown. We reveal that PZLS-1 is itself asymmetrically localised by Wnt-Ror-Dvl signalling, and is closely associated with DSH-1. PZLS-1 and DSH-1 colocalise in intracellular puncta and are mutually required for their localisation. *In vivo* structure-function analyses reveal conserved protein motifs that differentially control PZLS-1 localisation, abundance and function. This includes a previously uncharacterized highly conserved sequence that regulates PZLS-1 abundance in a DSH-1-dependent manner. We further show that PZLS-1 and DSH-1 control the abundance and polarised localisation of VAB-8/KIF26b, another conserved target of Wnt-Ror signalling. Together, these findings indicate that Wnt-Ror-Dvl signalling can control the spatial organisation and abundance of PDZRN- and KIF26B-family proteins *in vivo*, providing a mechanism to translate extracellular polarity cues into intracellular asymmetry.

## Introduction

Cell and tissue polarity can be established through mechanisms involving cell-cell communication or extrinsic morphogenetic cues like Wnt ligands. Wnt ligands are glycoproteins that function as short- or long-range signalling molecules, with permissive or instructive roles^1–3^. In metazoans, Wnt ligands contribute to polarisation processes such as the establishment of embryonic dorsoventral, anteroposterior and left-right axes, convergence and extension movements, asymmetric cell division or node cell polarisation. In some instances, the polarisation information is provided by gradients of extracellular Wnt proteins^4–6^. These Wnt gradients have been visualised directly in the *Drosophila* wing imaginal discs^7^ or *in vivo* in *C. elegans*^8,9^.

Wnt ligands can activate a diverse range of signalling pathways through different combinations of receptors and intracellular effectors^10^. These pathways are broadly classified into canonical Wnt/β-catenin signalling and non-canonical Wnt pathways. Among the latter, Wnt ligands can signal through Ror receptor tyrosine kinases, which can act in conjunction with Frizzled receptors^11^. In vertebrates, mice defective for Wnt5a or for Ror1 and Ror2 have similar phenotypes, suggesting that Wnt5a is the main ligand of Ror receptors *in vivo*^12–15^. Consistently, mutations in Wnt5a, Ror2, Frizzled2 and Dishevelled 1, 2 and 3 have been identified in cohorts of patients with Robinow syndrome, underscoring their crucial roles during development and tissue morphogenesis^16^.

A diverse set of effectors has been reported to act downstream of Wnt-Ror signalling, including Dishevelled^17^, Rac^18^, FLNa^17,19^, RhoA, and Daam1^20,21^. Recently, PDZRN3 and KIF26B have been identified as additional targets of Wnt-Ror-Dvl signalling. Indeed, proteomic screens showed that Wnt5a-Ror signalling targets both PDZRN3 and KIF26B for proteasomal degradation^22,23^. Moreover, both PDZRN3 and KIF26B interact with Dvl and participate in polarisation processes, in the retinal vasculature or in endothelial cell polarisation, respectively^24,25^.

PDZRN3/LNX3 belongs to the evolutionarily conserved LNX/PDZRN protein family, which also includes the paralogs PDZRN4/LNX4 and PDZD4/PDZRN4L/LNX5. A defining feature of these proteins is the C-terminal LNX3H domain, a domain of unknown function that is not found in any other metazoan protein. The LNX3H domain is evolutionarily ancient, with orthologs present in diverse metazoan lineages from cnidarians to vertebrates^26^. In vertebrates, it is associated with different combinations of PDZ (PSD-95, Discs large, and ZO-1) and RF (RING finger) domains, resulting in proteins with distinct domain architectures and suggesting functional diversification within the family.

PDZRN3 is the most studied member of the family, and its longest isoform contains a RING finger domain. PDZRN3 has therefore been studied primarily as an E3 ubiquitin ligase. However, ubiquitin ligase activity is unlikely to account for all functions of LNX3H-containing proteins. All PDZD4 isoforms lack a RING finger domain, as do shorter isoforms of PDZRN3 and PDZRN4. Consistent with this idea, a recent study uncovered the relevance of a short PDZRN4 isoform at the neuromuscular junction^27^. These observations suggest that LNX/Pdzrn proteins have important RING finger-independent roles, and therefore potentially ubiquitin ligase-independent functions.

Vertebrate PDZRN3 participates in a broad range of developmental processes, including myogenic differentiation and myotube formation^28–30^, kidney development^31^, nervous-system development^32^, neuromuscular-junction formation^33^, and retinal vascularization^24^. Its functions are not restricted to development: PDZRN3 also contributes to adult tissue homeostasis, including the maintenance of muscle mass, endothelial cell-cell junctions and vascular permeability, and the regulation of inflammatory and apoptotic responses^34–39^. Finally, altered *PDZRN3, PDZRN4* and *PDZD4* expressions have been reported in several cancers, where these genes have been implicated in tumour cell proliferation and dissemination^40–44^. In comparison with PDZRN3, however, PDZRN4 and PDZD4 remain poorly characterized, and little is known about the functions of these proteins or the roles of their conserved domains.

Recently, we discovered that a non-canonical Wnt signalling cascade involving EGL-20/Wnt, CAM-1/Ror and DSH-1/Dishevelled controls the formation of polarised membrane compartments at the plasma membrane of *C. elegans* muscle cells^45^. In the worm, 95 mononucleated striated muscle cells are distributed along the length of the animal. They are organised into four longitudinal quadrants, each made of two rows of intercalated cells^46^. We have shown that several proteins, including ion channels, membrane proteins and members of the dystrophin-associated protein complex, are asymmetrically distributed in these body wall muscles. For instance, the potassium channel TWK-28 is enriched at the anterior tip of each muscle cell. This asymmetric organisation is repeated in all cells along the anteroposterior axis of the animal, creating a remarkable tissue-wide pattern reminiscent of planar cell polarity. However, how the Wnt pathway implements polarity downstream of DSH-1 remained unknown.

In this study, we identify PZLS-1, the sole *C. elegans* member of the PDZRN family, as a regulator of muscle cell polarity. By endogenously tagging the protein, we revealed that PZLS-1 itself displays a polarised subcellular distribution, and that its asymmetric distribution depends on Wnt-Ror signalling. Furthermore, PZLS-1 colocalises with DSH-1 in intracellular puncta, and their localisation is mutually dependent. Using CRISPR/*Cas9* gene editing, we performed an *in vivo* structure-function analysis of PZLS-1. We identify conserved sequences required for PZLS-1 localisation and function, and uncover a novel, highly conserved motif that regulates PZLS-1 abundance downstream of DSH-1. Finally, we show that PZLS-1 and DSH-1 regulate both the abundance and subcellular localisation of VAB-8/KIF26. Conserved C-terminal regions of KIF26B that mediate Wnt5a-induced degradation also regulate VAB-8 abundance and localisation, revealing a conserved regulatory logic within the pathway. Together, these findings establish *C. elegans* muscle polarity as a powerful *in vivo* system for dissecting the molecular mechanisms linking Wnt-Ror signalling to PDZRN- and KIF26-family proteins.

## Results

### A forward genetic screen identifies the LNX3H-domain protein PZLS-1 as a regulator of muscle polarity

We previously showed that *C. elegans* body wall muscles (BWM) display a clear anteroposterior polarity that depends on a non-canonical Wnt signalling cascade involving EGL-20/Wnt, CAM-1/Ror and DSH-1/Dishevelled^45^. To identify additional regulators of muscle polarity acting downstream of this cascade, we performed a visual forward genetic screen, using the distribution of SGCB-1/β-Sarcoglycan as a polarity readout. In wild-type animals, SGCB-1 is enriched at the anterior tip of each muscle cell, where it forms a characteristic “comet-shaped” pattern, while regularly spaced SGCB-1 clusters occupy the remainder of the cell (Fig. 1A)^45^. We isolated three independent mutant lines in which this asymmetric distribution was disrupted, and in which the gene *C50F7.6* was mutated (Fig. S1A). These comprised two nonsense mutations (Gln254* and Trp286*) and one missense mutation (Pro321Leu) (Fig. 1C).

**Figure 1.**
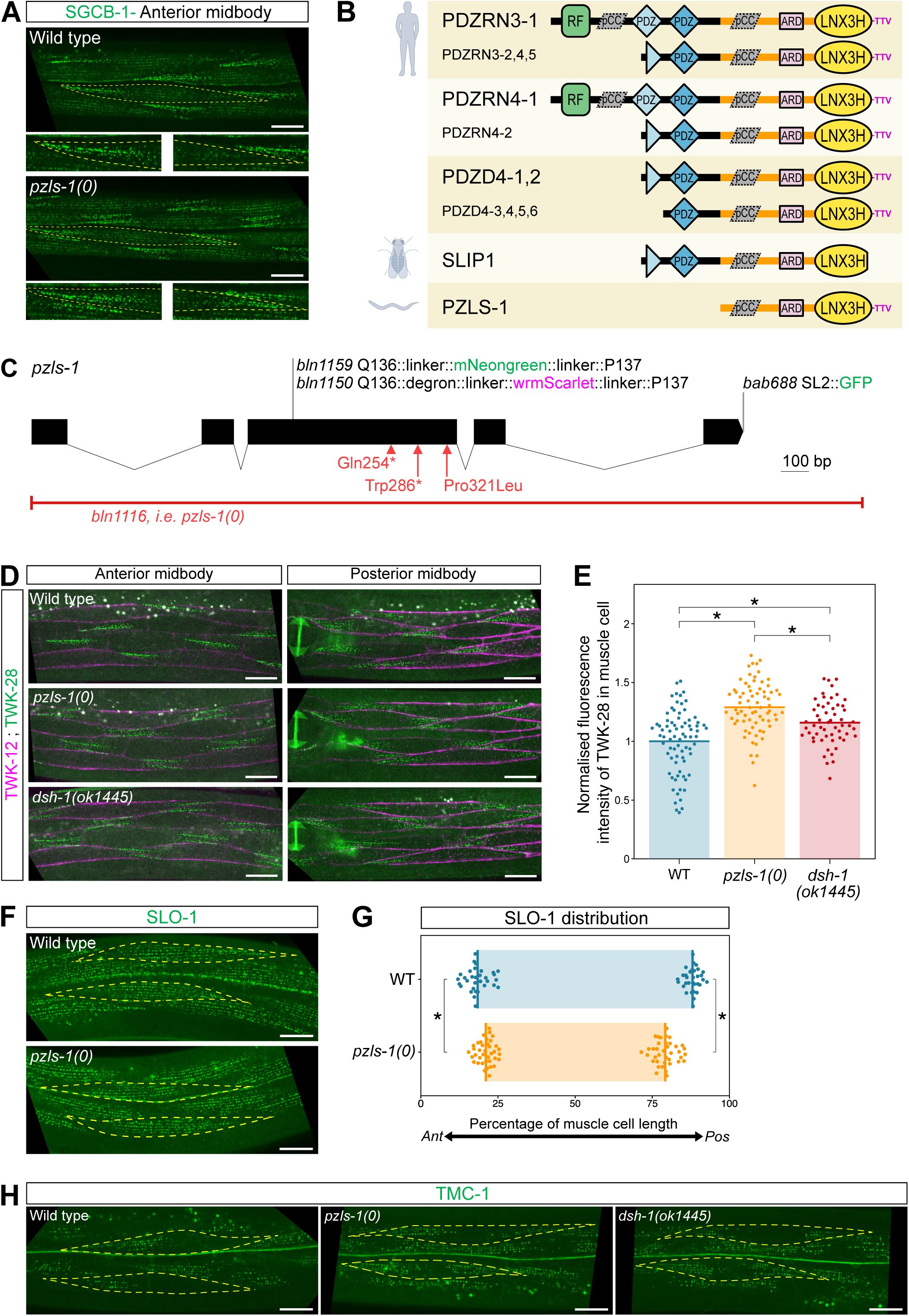
PZLS-1/PDZRN is a regulator of muscle polarity. **(A)** Asymmetrical distribution of SGCB-1/β-Sarcoglycan in a wild-type context *versus* symmetrical localisation in a *pzls-1* loss-of-function mutant. **(B)** Domain structure of LNX3H-containing proteins in human, *Drosophila* and *C. elegans*. RF: Ring Finger, pCC: predicted coiled-coil, ARD: abundance regulatory domain, TTV: C-terminal PDZ-binding motif. See *Methods* for accessions numbers of corresponding human isoforms. **(C)** *pzls-1* genomic locus. Mutant alleles, in red. *pzls-1* molecular null allele, *bln1116*, hereafter referred to as *pzls-1(0)*. Translational fusion reporters, *bln1150* and *bln1159.* Transcriptional reporter, *bab688.* **(D)** Symmetrical distribution of TWK-28 in *dsh-1(0)* loss-of-function and *pzls-1(0)* backgrounds, in muscles of the anterior or posterior midbody. Muscle membranes are labelled by TWK-12-wrmScarlet (in magenta). **(E)** Quantification of total mNeonGreen-TWK-28 fluorescence. Normalized fluorescence intensity relative to mean fluorescence of WT. Each data point represents one muscle cell; bars represent data means; *n =* 77, 72, 56 (for respective columns); *N* = 3 independent experiments; Kruskal-Wallis followed by Dunn’s test with Holm’s correction. * p < 0.01. **(F)** Symmetrical distribution of SLO-1 in a *pzls-1(0)* loss-of-function background. **(G)** Quantification of SLO-1 domain boundary positions in wild type and *pzls-1(0)*. Coloured bars indicate the region containing SLO-1. Bars represent data means; Number of animals assayed: *n =* 32, 38 (for respective lines); *N* = 2 independent experiments; Welch t-test, * p < 0.01. **(H)** Asymmetrical distribution of TMC-1 in body wall muscles in a wild-type context, and symmetrical distribution of TMC-1 in *dsh-1(0)* and *pzls-1(0)* loss-of-function backgrounds. Muscle cell outlines are indicated with yellow dashed lines. Anterior to the left. Scale bars, 20 µm. Raw data and statistical analyses are presented in the Source Data file.

To establish that loss of *C50F7.6* was responsible for the polarity phenotype, we generated a full deletion of the locus by CRISPR/*Cas9* gene editing (Fig. 1C). In *C50F7.6* mutants, SGCB-1 accumulated at both extremities of the cell, resulting in a symmetrical distribution (Fig. 1A). We previously showed that EGL-20/Wnt and CAM-1/Ror are predominantly required for polarity in posterior midbody muscles, whereas DSH-1 is required in muscles both anterior and posterior to the vulva^45^. Similar to *dsh-1* mutants, loss of *C50F7.6* disrupted SGCB-1 polarity on both sides of the vulva (Fig. S1B), indicating that *C50F7.6* is broadly required for SGCB-1 asymmetry across the midbody musculature.

Sequence comparison identified C50F7.6 as the sole *C. elegans* member of the LNX3H-containing branch of the PDZRN/LNX protein family. C50F7.6 contains a characteristic C-terminal LNX3H domain which is also present in PDZRN3, PDZRN4 and PDZD4 (also named LNX3, LNX4 and LNX5, respectively)^26^ (Fig. 1B and S1E). *Drosophila* similarly encodes a single related protein, Slip1, which contains a partially truncated LNX3H domain^26^. Sequence comparison further revealed conserved regions N-terminal to the LNX3H domain, including a predicted coiled-coil region and a short motif containing a highly conserved “SSAYNT” signature sequence (Fig. 1B and S1E). We therefore named C50F7.6 “PZLS-1”, for “<u>P</u>D<u>Z</u>RN3, PDZRN4, <u>L</u>NX5 and <u>S</u>lip1 ortholog 1”. Unlike the longest vertebrate PDZRN3 and PDZRN4 isoforms, PZLS-1 lacks an N-terminal RING finger domain, as do PDZD4, Slip1 and several vertebrate PDZRN3 and PDZRN4 isoforms (Fig. 1B).

We next asked whether loss of PZLS-1 affected other polarised membrane proteins. We previously showed that loss of *dsh-1* converts the asymmetric comet-shaped distribution of the potassium channel TWK-28 into a symmetrical pattern, with TWK-28 accumulating at both extremities of the muscle cell^45^. Indeed, loss of PZLS-1 produced a similar phenotype as TWK-28 was now present both at the anterior and the posterior end of muscle cells (Fig. 1D and S1C). Using *in vivo* confocal imaging, we estimated that total TWK-28-associated fluorescence was modestly increased in both mutants relative to wild type (Fig. 1E and S1D), but this increase was substantially smaller than would be expected from a doubling of the TWK-28 signal. Thus, except for a small difference in TWK-28 abundance, PZLS-1 and DSH-1 affect TWK-28 polarity in a similar fashion.

Next, we analysed the distribution of the BK potassium channel SLO-1, which provided a distinct polarity readout as it is excluded from the anterior portion of each muscle cell and covers most of the remainder of the cell surface in a punctate pattern (Fig. 1F)^45^. In *pzls-1* mutants, SLO-1 was absent from both extremities and distributed in a symmetrical pattern (Fig. 1F, G).

Finally, we examined TMC-1, a mechanosensitive ion channel, which appeared to have an asymmetrical distribution in head muscles according to previously published data^47,48^. We confirmed that TMC-1 displayed a polarised distribution with dispersed puncta preferentially occupying the anterior region of the cell (Fig. 1H). This pattern was morphologically distinct from the compact anterior enrichment of TWK-28. Nevertheless, TMC-1 polarity also depended on both DSH-1 and PZLS-1. Indeed, loss of either protein resulted in a symmetrical distribution, with TMC-1 detected at both ends of the muscle cell (Fig. 1H).

Thus, PZLS-1 is required for the asymmetric distribution of membrane proteins with several distinct spatial patterns, including SGCB-1, TWK-28, SLO-1 and TMC-1. Moreover, loss of *pzls-1* consistently produced polarity defects similar to *dsh-1* mutants. Together, these results identify PZLS-1 as a novel regulator of muscle cell polarity.

### PZLS-1 functions in muscle and is asymmetrically localised

To determine the tissues in which PZLS-1 is expressed, we first examined available transcriptomic data and generated a transcriptional knock-in reporter line (Fig. 1C). In the CeNGEN adult hermaphrodite dataset, *pzls-1* expression is highest in body wall muscles (BWMs), vulval muscles and a subset of neurons (Fig. S2A). Consistent with these data, our transcriptional reporter was expressed in several muscle types, including BWMs, vulval muscles and the anal depressor muscle, as well as in neurons (Fig. S2B).

We next generated endogenous PZLS-1 fusions with either wrmScarlet or mNeonGreen to examine its subcellular distribution in BWMs (Fig. 1C, 2A, B). Strikingly, PZLS-1 was asymmetrically distributed along the anteroposterior axis of muscle cells, with a pronounced enrichment at the posterior end (Fig. 2A). Thus, PZLS-1 is not only required for muscle polarity but is itself polarised within muscle cells.

**Figure 2.**
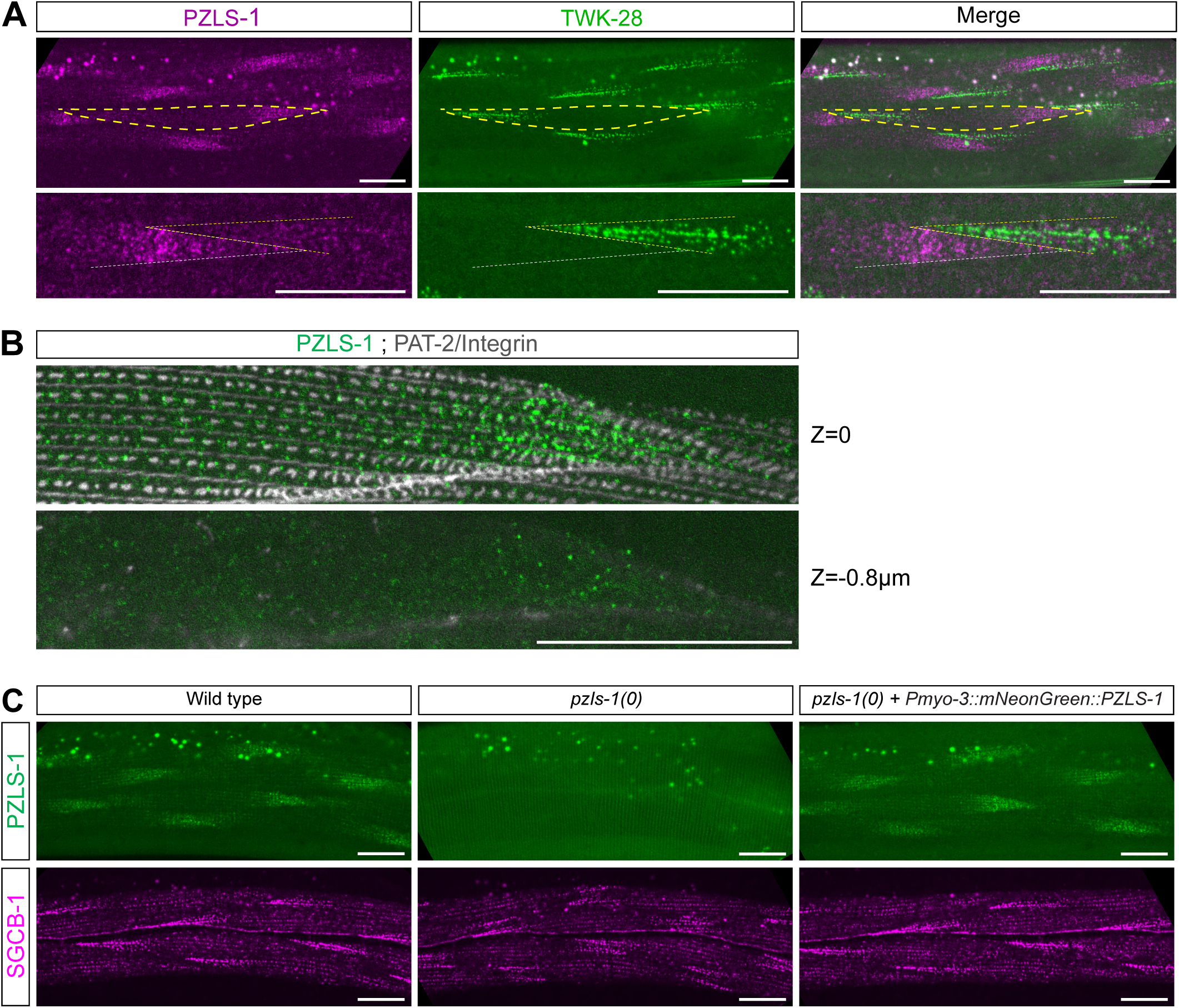
PZLS-1 is posteriorly enriched and cell-autonomously required in muscle cells. **(A)** PZLS-1 and TWK-28 are enriched at opposite ends of muscle cells. View of anterior midbody region (*first row*). Magnified view of contact site between two consecutive muscle cells (*second row*). Muscle cell outlines are indicated with yellow or white dashed lines. **(B)** PZLS-1 (green) is enriched at or close to the cell surface, based on PAT-2/Integrin labelling (white) (*first row*). PZLS-1 puncta are also found inside the muscle cell (*second row*). **(C)** Muscle-specific expression of a mNeonGreen-PZLS-1 cDNA rescues SGCB-1 localisation in *pzls-1(0)*. Anterior to the left. Scale bars, 20 µm.

Closer examination revealed several distinct subcellular pools of PZLS-1 (Fig. 2B). To relate its subcellular localisation to muscle architecture, we compared it with PAT-2/integrin, which labels dense bodies and M-lines^46^. In the central region of muscle cells, PZLS-1 formed small puncta close to the cell surface. These puncta were located between or adjacent to dense bodies, but did not colocalise with them. Towards the posterior end of the cell, surface-associated PZLS-1 puncta formed a denser pattern, forming the prominent posterior enrichment. In addition, a population of intracellular PZLS-1 puncta was detected in the posterior region (Fig. 2B). These observations indicate that PZLS-1 occupies several subcellular compartments and that both its surface-associated and intracellular pools are spatially polarised.

Because PZLS-1 is expressed in BWMs, we next asked whether it acts cell-autonomously to establish muscle polarity. As described above, loss of *pzls-1* disrupts the asymmetric distribution of SGCB-1/β-Sarcoglycan. Expression of a *pzls-1* cDNA specifically in BWMs was sufficient to restore SGCB-1 asymmetry (Fig. 2C), demonstrating that PZLS-1 can act within muscle cells to control their polarity. Independent transgenic lines expressing different levels of PZLS-1 all restored SGCB-1 polarity (Fig. S2C), indicating that polarity can be rescued across a range of PZLS-1 expression levels.

Together, these results show that PZLS-1 acts cell-autonomously in BWMs and is itself asymmetrically organised into distinct subcellular pools, with a prominent enrichment at the posterior end of muscle cells.

### PZLS-1 localisation is controlled by Wnt-Ror-Dvl signalling and PZLS-1 colocalises with DSH-1 in intracellular puncta

In vertebrates, PDZRN3 has been functionally linked to Wnt-Ror-Dvl signalling^23,24^. Because EGL-20/Wnt, CAM-1/Ror and DSH-1/Dvl are required for muscle polarity in *C. elegans*^45^, we asked whether these proteins also control PZLS-1 localisation.

In posterior midbody BWMs, loss of either EGL-20 or CAM-1 disrupted the posterior enrichment of PZLS-1 (Fig. 3A), in the same region in which these mutants cause polarity defects^45^. Loss of DSH-1 similarly abolished the posterior enrichment of PZLS-1, but affected muscle cells throughout the body. Moreover, in anterior midbody muscles, where intracellular PZLS-1 puncta were most readily observed, these puncta were no longer detected in *dsh-1* mutants (Fig. 3B). Thus, EGL-20/Wnt, CAM-1/Ror and DSH-1/Dvl are required for the polarised localisation of PZLS-1.

**Figure 3.**
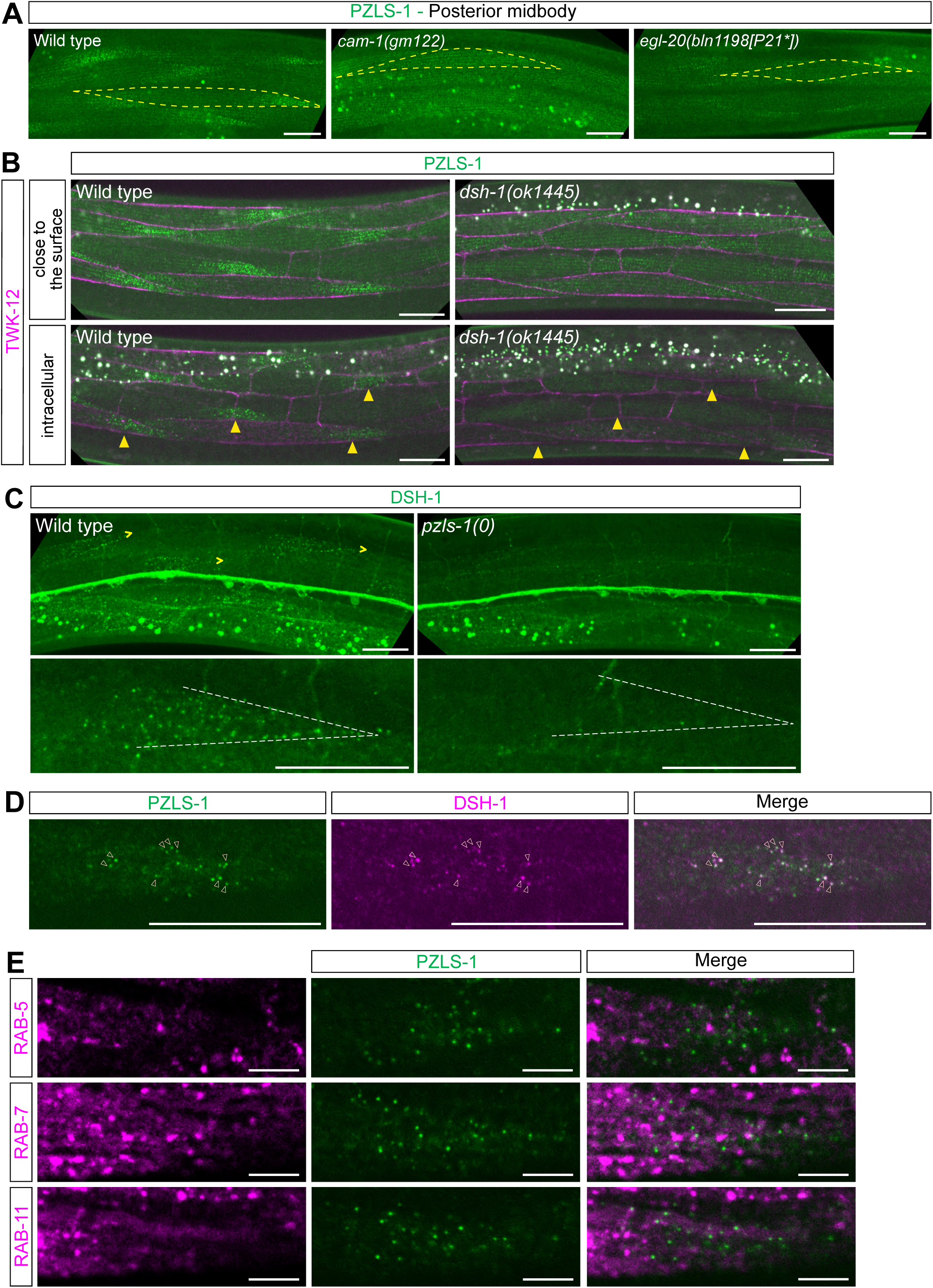
Asymmetric localisation of PZLS-1 depends on Wnt-Ror signalling and is mutually dependent on DSH-1. **(A)** CAM-1/Ror and EGL-20/Wnt are required for the asymmetric localisation of PZLS-1. Muscle cell outlines are indicated with yellow dashed lines. **(B)** DSH-1 is required for posterior enrichment of PZLS-1. Yellow arrowheads indicate posterior end of muscle cells. *Top row,* surface of muscle cells. *Bottom row*, interior of muscle cells. Muscle membranes are labelled by TWK-12-wrmScarlet (in magenta). **(C)** PZLS-1 is required for the asymmetrical localisation of DSH-1. *First row*, right-pointing yellow arrowheads indicate posterior extremity of muscle cells. *Second row*, magnified view of the posterior end of a muscle cell. Muscle cell outlines are indicated with white dashed lines. **(D)** PZLS-1 and DSH-1 colocalise in intracellular puncta at the posterior end of muscle cells. Open arrowheads label colocalised puncta. **(E)** Intracellular PZLS-1 puncta do not colocalise with RAB-5, RAB-7 or RAB-11 in body wall muscle cells. Anterior to the left. Scale bars, 20 µm.

We next asked whether PZLS-1 reciprocally influences DSH-1 localisation. DSH-1 itself displays an asymmetric distribution in BWMs, including intracellular puncta enriched in the posterior end of anterior midbody muscles^45^. Strikingly, these polarised DSH-1 puncta were lost in *pzls-1* mutants (Fig. 3C). PZLS-1 and DSH-1 are therefore mutually dependent for their subcellular localisation.

The similar distribution of PZLS-1 and DSH-1 puncta prompted us to test whether the two proteins occupy the same intracellular compartments. Indeed, PZLS-1 and DSH-1 partially colocalised within posterior intracellular puncta (Fig. 3D). We next asked whether these structures corresponded to endosomal compartments. This possibility was of particular interest because Wnt5a induces internalisation of FZD4 and DVL3 into early endosomes in HeLa cells in a PDZRN3-dependent manner^24^, although DVL proteins can also form intracellular puncta independently of endosomal compartments^49–51^. In BWMs, however, PZLS-1 and DSH-1 puncta did not colocalise with either RAB-5-, RAB-7- or RAB-11-labelled compartments (Fig. 3E), which correspond to early, late or recycling endosomes, respectively.

Interestingly, a subset of DSH-1 puncta partially colocalised with EGL-20 (Fig. S3A). EGL-20 is secreted from cells located posteriorly in the animal and is not detectably expressed in the muscle cells in which these puncta were observed^52^. This observation raises the possibility that at least some DSH-1/PZLS-1-positive puncta contain extracellularly derived EGL-20 and may represent intracellular signalling assemblies. Their precise identity, however, remains to be determined.

Taken together, these results place PZLS-1 localisation downstream of EGL-20/CAM-1/DSH-1 signalling and reveal a reciprocal requirement for PZLS-1 and DSH-1 for their polarised localisation. PZLS-1 and DSH-1 further colocalise in a subset of intracellular puncta that are distinct from the canonical endosomal compartments examined here.

### Conserved LNX3H residues are required for PZLS-1 function downstream of DSH-1-dependent localisation

In vertebrates, Wnt5a-Ror signalling leads to phosphorylation of PDZRN3 at three residues within the LNX3H domain, promoting PDZRN3 degradation. Perturbation of these phosphorylation sites impairs cell migration^23^. These residues are conserved in human PDZRN4 and PDZD4/LNX5, as well as in *C. elegans* PZLS-1 (Fig. 4A). We therefore asked whether the corresponding residues are required for PZLS-1 function.

**Figure 4.**
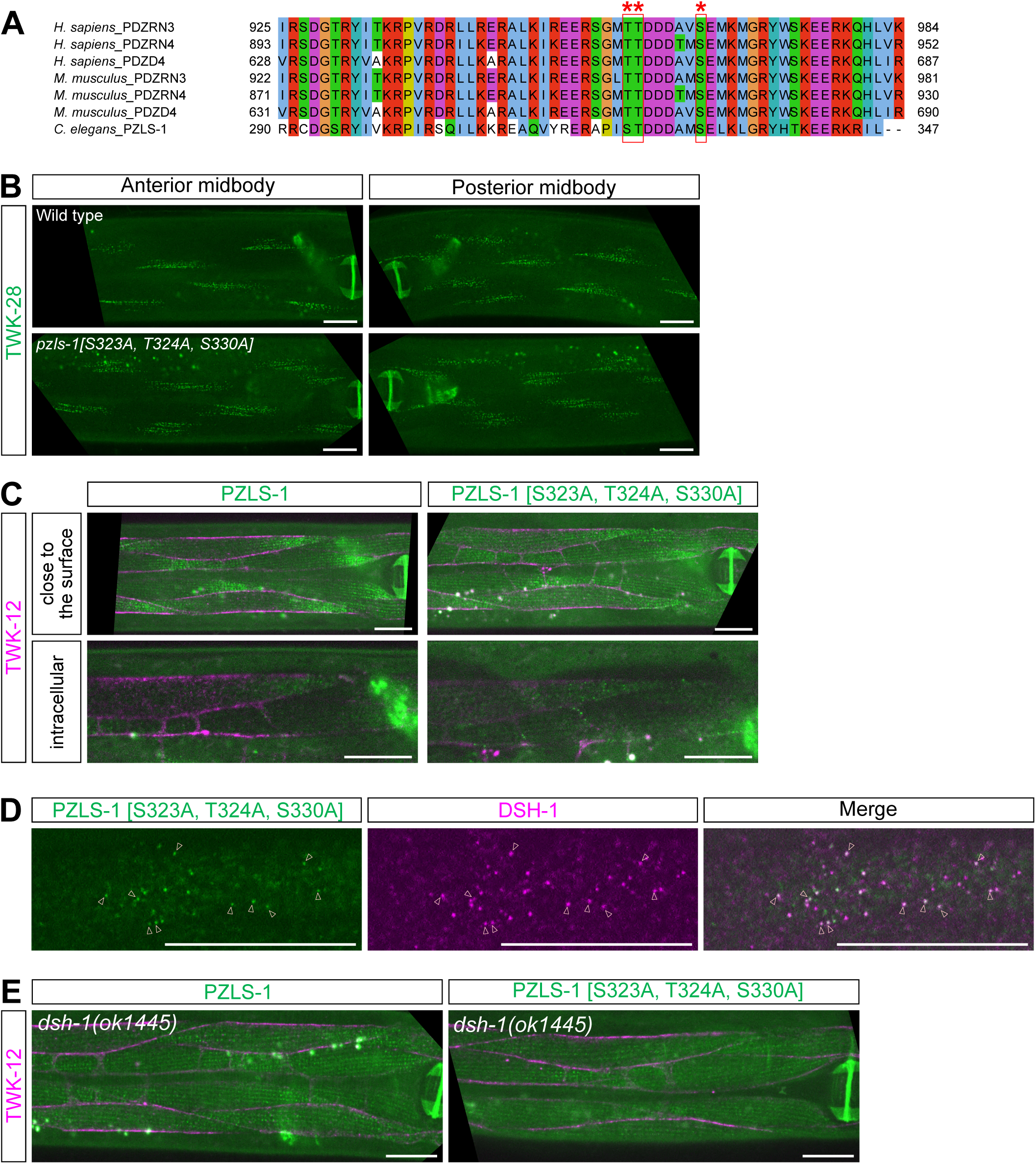
Conserved putative phosphorylation sites are required for PZLS-1 function but largely dispensable for its localisation. **(A)** Sequence conservation of putative PDZRN phosphorylation sites in human, mouse and *C. elegans* LNX3H proteins. Residues are labelled by red asterisks and boxes. **(B)** Asymmetric localisation of TWK-28 is lost in a PZLS-1([S323A, T324A, S330A]) triple mutant. **(C)** Posterior PZLS-1 enrichment is only partly affected upon mutation of all three S/T residues. **(D)** Intracellular colocalisation of PZLS-1 and DSH-1 is preserved in a PZLS-1([S323A, T324A, S330A]) triple mutant. Open arrowheads indicate colocalised puncta. **(E)** Posterior enrichment of PZLS-1 and PZLS-1([S323A, T324A, S330A]) requires DSH-1. Muscle membranes are labelled by TWK-12-wrmScarlet in panels *C* and *E* (in magenta). Anterior to the left. Scale bars, 20 µm.

We simultaneously substituted the three conserved residues with alanines to generate a putative phosphorylation-deficient PZLS-1 variant. These mutations caused a strong loss-of-function phenotype: TWK-28 polarity was disrupted in muscles both anterior and posterior to the vulva, closely resembling the phenotype of *pzls-1* null mutants (Fig. 1D and 4B). Thus, these conserved LNX3H residues are essential for PZLS-1-dependent muscle polarity.

We next asked whether this resulted from altered PZLS-1 stability or localisation. We thus introduced the same substitutions into an endogenously mNeonGreen-tagged PZLS-1 reporter line. The mutant protein remained readily detectable and retained an asymmetric distribution (Fig. 4C). Although its posterior enrichment appeared somewhat reduced compared with wild-type PZLS-1, intracellular puncta remained clearly visible and continued to colocalise with DSH-1 (Fig. 4D). Thus, mutation of these residues strongly impairs PZLS-1 function while only partially affecting its subcellular distribution, arguing against a loss of protein stability or gross mislocalisation as the primary cause of the phenotype.

We then asked how these residues relate to the DSH-1-dependent localisation of PZLS-1. In a *dsh-1* null background, both wild-type PZLS-1 and the alanine variant lost their polarised distribution and intracellular puncta, becoming essentially uniformly distributed within muscle cells (Fig. 4E). Thus, because the triple mutant protein can still be recruited to polarised intracellular puncta in the presence of DSH-1 but is non-functional, these residues appear to be required for PZLS-1 activity after its DSH-1-dependent localisation.

Together, these results identify three conserved residues within the LNX3H domain that are essential for PZLS-1 function but largely dispensable for its polarised localisation, separating DSH-1-dependent recruitment of PZLS-1 from its subsequent activity in establishing muscle polarity.

### A conserved PDZ binding domain is only partly required for PZLS-1 localisation and function

PZLS-1 and DSH-1 colocalise in intracellular puncta, and their vertebrate orthologs have been reported to interact through the PDZ domain of DVL3 and the C-terminal PDZ-binding motif of PDZRN3^24^. Both of these domains are conserved in the corresponding *C. elegans* proteins. We therefore asked whether this putative interaction contributes to the reciprocal localisation and function of DSH-1 and PZLS-1.

We previously showed that a DSH-1 protein lacking its PDZ domain fails to rescue the polarity defects of *dsh-1* null mutants^45^. Consistently, deletion of the PDZ domain of DSH-1 by CRISPR/*Cas9* gene editing disrupted TWK-28 polarity (Fig. S4A). To determine whether the PDZ domain also controls DSH-1 and PZLS-1 localisation, we deleted it in an endogenously tagged DSH-1 reporter line. The truncated protein remained readily detectable in neurons in the ventral nerve cord (Fig. 5A), arguing against gross destabilisation of DSH-1. In muscle cells, however, the intracellular DSH-1 puncta that depend on PZLS-1 were lost. Conversely, PZLS-1 localisation was also disrupted, as it was no longer asymmetrically distributed. Thus, deletion of the DSH-1 PDZ domain affects the polarised localisation of both proteins, consistent with this domain contributing to their functional association.

**Figure 5.**
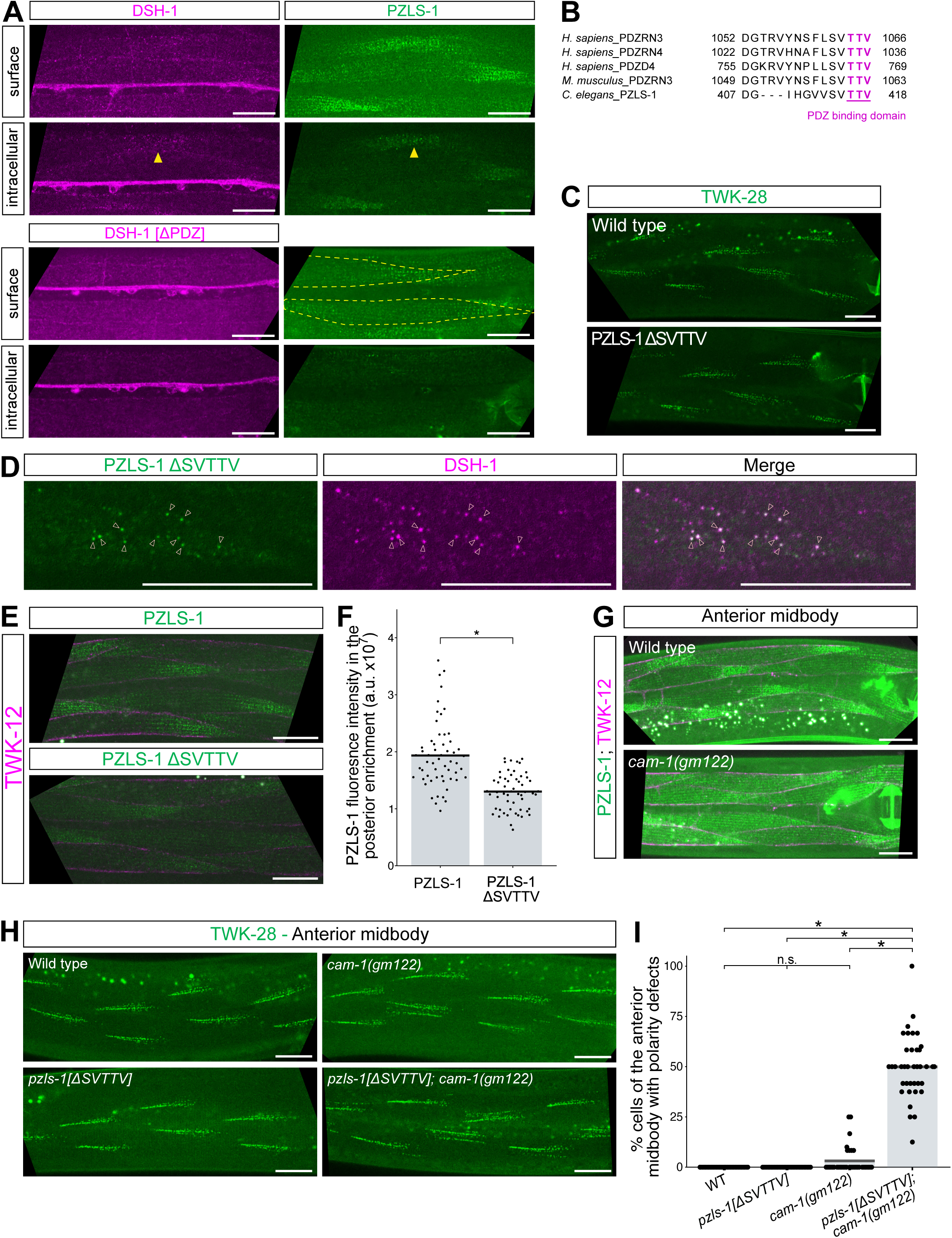
The DSH-1 PDZ domain is necessary for PZLS-1 localisation, whereas the PZLS-1 PDZ-binding motif is only partially required. **(A)** Deletion of the DSH-1 PDZ domain abolishes intracellular DSH-1 puncta and disrupts the posterior enrichment of PZLS-1. **(B)** The PDZ-binding motif of vertebrate LNX3H proteins is conserved in PZLS-1. **(C)** Deletion of the PZLS-1 PDZ-binding motif does not affect TWK-28 polarity. **(D)** Deletion of the PZLS-1 PDZ-binding motif does not prevent PZLS-1 from colocalising with DSH-1. Open arrowheads indicate colocalised puncta. **(E)** Deletion of the PZLS-1 PDZ-binding motif does not abolish its posterior enrichment but reduces PZLS-1 abundance within this domain. **(F)** Quantification of PZLS-1-mNeonGreen fluorescence intensity. Each data point represents one muscle cell; bars represent data means; *n =* 56, 56 (for respective columns); *N* = 2 independent experiments; Two-tailed Mann-Whitney U test. *p < 0.01. **(G)** Loss of CAM-1/Ror affects the posterior enrichment of PZLS-1 in anterior midbody muscles. **(H)** Combined *cam-1* loss-of-function and PZLS-1 PDZ-binding motif deletion causes synthetic polarity defects. **(I)** Quantification of polarity defects in the anterior midbody, in the muscle cells number 9 to 14 of muscle quadrants. Polarity defects were defined as the presence of a consistent signal in the posterior tip of the cell, being at least 2/3 of the anterior comet in length. Each data point represents one muscle cell; bars represent data means; *n =* 33, 32, 35, 38 (for respective columns); *N* = 2 independent experiments; Kruskal-Wallis followed by Dunn’s test with Holm’s correction. n.s., not significant, * p<0.01. Muscle membranes are labelled by TWK-12-wrmScarlet in panels *E* and *G* (in magenta). Anterior to the left. Scale bars, 20 µm. Raw data and statistical analyses are presented in the Source Data file.

We next deleted the conserved PDZ-binding motif of PZLS-1 (Fig. 5B), expecting that this might produce the same phenotype. Surprisingly, deletion of this motif did not disrupt TWK-28 polarity (Fig. 5C). The truncated PZLS-1 protein also retained its asymmetric localisation and continued to colocalise with DSH-1 in intracellular puncta (Fig. 5D, E). Thus, the PDZ-binding motif appeared not to be strictly required *in vivo* for either PZLS-1 localisation or its role in establishing muscle polarity.

Nevertheless, quantitative analysis revealed that the amount of PZLS-1(ΔSVTTV) within the posterior enrichment was reduced compared with wild type PZLS-1 (Fig. 5E, F). This suggested that the motif contributes to efficient accumulation of PZLS-1 within this polarised domain. Under otherwise wild-type conditions, the residual pool of PZLS-1(ΔSVTTV) may therefore be sufficient to maintain polarity. We reasoned that its reduced accumulation might however become functionally limiting in a sensitised genetic background. We therefore tested PZLS-1(ΔSVTTV) function under conditions in which Wnt-Ror signalling was partially compromised.

Indeed, in anterior midbody muscles, loss of CAM-1 partially reduced the posterior enrichment of PZLS-1 (Fig. 5G), while TWK-28 polarity defects remained rare (Fig. 5H, I), indicating that polarity is largely maintained in the absence of CAM-1. Strikingly, combining *cam-1* loss-of-function with deletion of the PZLS-1 PDZ-binding motif markedly increased the frequency of TWK-28 polarity defects (Fig. 5H, I). Thus, although the PDZ-binding motif is dispensable for polarity in an otherwise wild-type background, its contribution becomes functionally important when Wnt-Ror signalling is compromised.

These genetic interactions also reveal a contribution of CAM-1 to the polarisation of anterior midbody muscles that is largely masked under wild-type conditions. The residual signalling present in *cam-1* mutants is sufficient to maintain TWK-28 polarity when PZLS-1 is intact, but becomes insufficient when PZLS-1 function is weakened by removal of its PDZ-binding motif. This suggests that CAM-1 contributes to anterior muscle polarity together with additional signalling inputs.

Together, these results show that the conserved PZLS-1 PDZ-binding motif is not absolutely required for PZLS-1 localisation or function, but promotes its efficient accumulation within the posterior polarised domain and becomes functionally important when Wnt-Ror signalling is reduced. They further support a role for the DSH-1 PDZ domain in coordinating the polarised localisation of DSH-1 and PZLS-1.

### A conserved β-sheet motif within the LNX3H domain is required for PZLS-1 localisation and function

The PDZ domain of DSH-1 is required for PZLS-1 localisation and muscle polarity, whereas the canonical PDZ-binding motif of PZLS-1 is only partially required. We thus considered the possibility that additional protein motifs might contribute to the association between the two proteins. To identify candidate interaction surfaces, we modelled a PZLS-1-DSH-1 heteromer using AlphaFold-Multimer (Fig. 6A and S5A). The predicted complex recapitulated the canonical interaction between the C-terminal PDZ-binding motif of PZLS-1 and the peptide-binding groove of the DSH-1 PDZ domain^53,54^. In addition, the model predicted a second interface between the N-terminal portion of the PZLS-1 LNX3H domain and the PDZ domain of DSH-1 (Fig. 6A). This sequence was proposed to adopt an extended β-sheet conformation^26^, and is likewise predicted by AlphaFold to form a similar structure in human PDZRN3, PDZRN4 and PDZD4/LNX5. Strikingly, the predicted interface involves five residues that are highly conserved across LNX3H-containing proteins (Fig. 6B). We therefore tested whether this conserved structural element contributes to PZLS-1 localisation and function.

**Figure 6.**
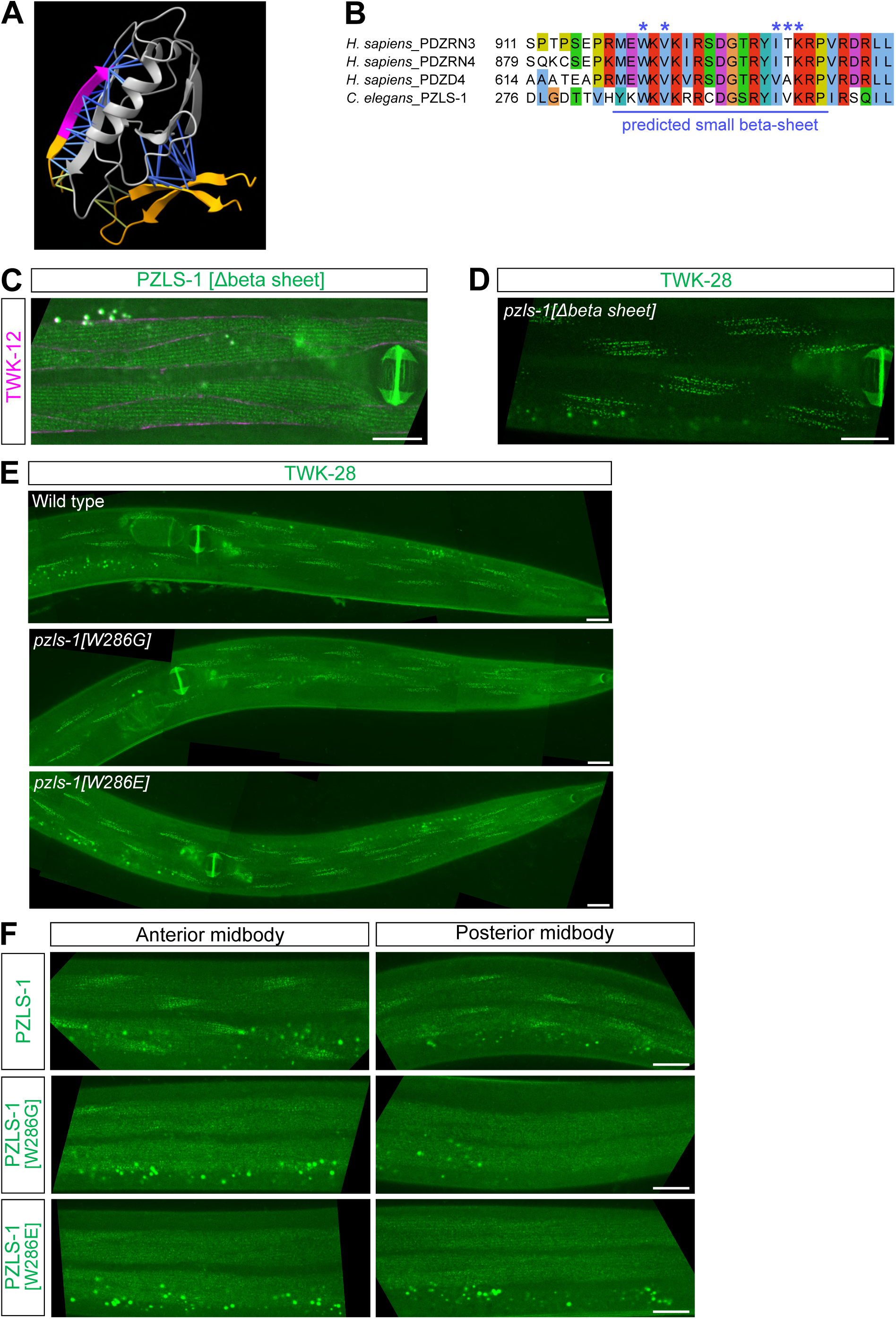
A conserved β-sheet within the LNX3H domain is required for PZLS-1 function and localisation. **(A)** Structural prediction of putative PZLS-1-DSH-1 protein-protein contacts (blue) using AlphaFold-Multimer. Only the PDZ domain of DSH-1 is shown (in grey), as well as portions of PZLS-1 predicted to interact with DSH-1 PDZ domain, including the PDZ binding motif (in magenta). **(B)** Sequence conservation of a predicted β-sheet structure within human and nematode LNX3H-containing proteins. Blue asterisks indicate residues within the LNX3H domain predicted to contact the PDZ domain of DSH-1 by AlphaFold-Multimer. **(C)** Deletion of the predicted β-sheet disrupts asymmetric localisation of PZLS-1. Muscle membranes are labelled by TWK-12-wrmScarlet (in magenta). **(D)** Deletion of the predicted β-sheet disrupts TWK-28 polarity. **(E)** W286G and W286E substitutions in PZLS-1 cause polarity defects in posterior midbody muscles. **(F)** W286G and W286E substitutions in PZLS-1 impair its posterior enrichment in midbody muscles. Anterior to the left. Scale bars, 20 µm.

Using CRISPR/*Cas9* gene editing, we replaced the predicted β-sheet at the endogenous *pzls-1* locus with a short flexible linker. This deletion/substitution caused a clear loss-of-function phenotype: TWK-28 polarity was disrupted and PZLS-1 lost its asymmetric distribution (Fig. 6C, D). These observations are consistent with a requirement for this conserved region in PZLS-1 localisation and signalling. However, because replacement of an entire β-sheet could perturb the local or global structure of PZLS-1, this experiment alone could not distinguish a specific functional requirement for the predicted interface from a more general structural defect.

To minimise potential effects on protein folding, we therefore introduced single amino-acid substitutions at conserved residues predicted to contribute to the interface. We focused on W286, which was predicted to make multiple contacts with residues on the face of the DSH-1 PDZ domain opposite the canonical PDZ-binding-motif interface. Substitution of W286 with either glutamic acid or glycine caused polarity defects in posterior midbody muscles (Fig. 6E). Both substitutions also reduced or abolished the posterior enrichment of PZLS-1 (Fig. 6F). Importantly, clear PZLS-1 enrichment remained detectable in the most anterior head muscles (Fig. S5B), indicating that these point mutations produce a weaker localisation defect than replacement of the entire β-sheet or complete loss of DSH-1 (Fig. S5C).

Together, these results identify a conserved β-sheet within the LNX3H domain that is required for PZLS-1 localisation and function. Its predicted association with the DSH-1 PDZ domain provides a potential explanation for why disruption of the canonical PZLS-1 PDZ-binding motif alone only partially impairs signalling. These findings suggest that multiple interaction surfaces may contribute to the functional association between PZLS-1 and DSH-1.

### A previously uncharacterized conserved sequence regulates PZLS-1 abundance

We next investigated the function of a highly conserved sequence located N-terminal to the LNX3H domain (Fig. 7A). This region was originally identified by Katoh and Katoh, who noted its conservation between PDZRN3, PDZRN4 and the *Drosophila* protein Slip1 and designated it as PR34H1. No known protein domain was detected within this sequence by RPS-BLAST, and its function had remained unexplored^55^. Sequence comparison revealed that this region is also conserved in *C. elegans* PZLS-1, including a highly conserved SSAYNT core followed by ESCRS residues that are shared between *C. elegans* and vertebrate proteins (Fig. 7A and S1E).

**Figure 7.**
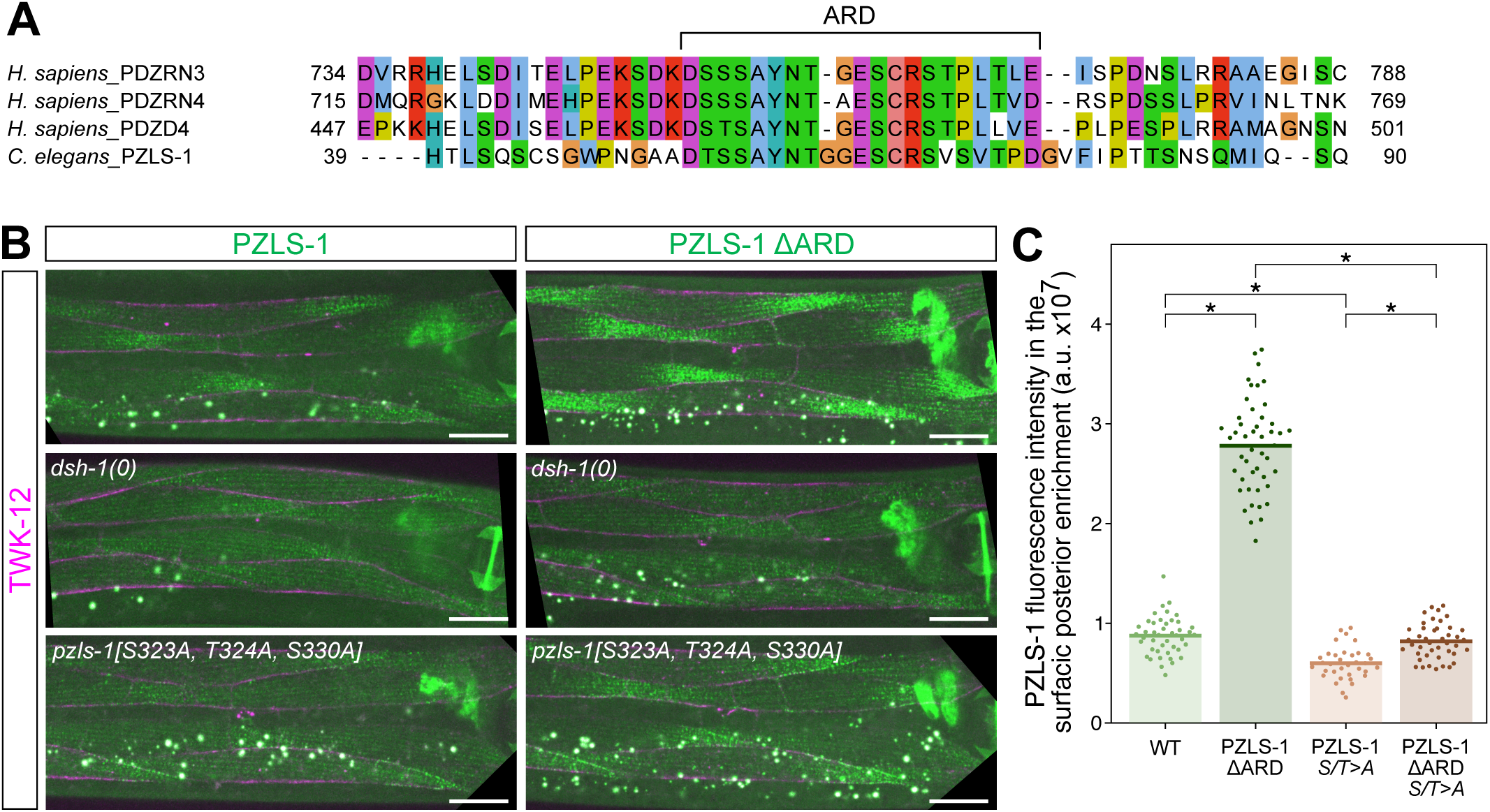
A conserved motif regulates PZLS-1 abundance *in vivo*. **(A)** Conservation of the abundance regulatory domain (ARD) sequence in human and *C. elegans* LNX3H-domain proteins. **(B)** Deletion of the ARD increases mNeonGreen-PZLS-1 fluorescence. This increase is abolished in a *dsh-1* loss-of-function mutant and markedly reduced in a PZLS-1([S323A, T324A, S330A]) triple mutant. Muscle membranes are labelled by TWK-12-wrmScarlet (in magenta). Anterior to the left. Scale bars, 20 µm. **(C)** Quantification of mNeonGreen-PZLS-1 fluorescence in wild-type PZLS-1 and PZLS-1([S323A, T324A, S330A]) triple mutant, with or without the ARD motif. Each data point represents one muscle cell; bars represent data means; *n =* 44, 35, 47, 43 (for respective columns); *N* = 2 independent experiments; Kruskal-Wallis test followed by Dunn’s test with Holm’s correction. * p<0.01. Raw data and statistical analyses are presented in the Source Data file.

To investigate its function, we used CRISPR/*Cas9* gene editing to delete this conserved sequence from a fluorescently tagged PZLS-1 knock-in reporter. Remarkably, this deletion increased PZLS-1 fluorescence (Fig. 7B, C), indicating that the sequence normally acts to limit PZLS-1 abundance. We therefore termed this conserved region the “<u>A</u>bundance <u>R</u>egulatory <u>D</u>omain” (ARD).

In vertebrates, Wnt5a-Ror-Dvl signalling promotes PDZRN3 degradation^23^. Although our genetic data argue against a simple model in which EGL-20-CAM-1 signalling establishes muscle polarity primarily by depleting PZLS-1, the strong increase in PZLS-1-associated fluorescence following ARD deletion suggested that its protein levels are nevertheless actively regulated. We therefore asked whether this regulation depends on DSH-1. In a *dsh-1* loss-of-function background, ARD deletion no longer caused the pronounced increase in PZLS-1 fluorescence observed in an otherwise wild-type background (Fig. 7B). Thus, the effect of the ARD on PZLS-1 abundance depends on DSH-1.

We next examined whether the three conserved LNX3H residues previously implicated in vertebrate PDZRN3 phosphorylation also contribute to ARD-dependent regulation. As shown above, these residues are conserved in PZLS-1 and are required for its function (Fig. 4). When all three residues were replaced by alanines, ARD deletion produced only a modest increase in PZLS-1 fluorescence. Quantification of fluorescence intensity showed that ARD deletion increased PZLS-1 levels by more than threefold in the wild-type protein, but only approximately 1.4-fold in the triple mutant variant (Fig. 7C). Thus, most of the increase in PZLS-1 abundance caused by ARD deletion depends on these conserved LNX3H residues.

Taken together, these results identify a previously uncharacterized conserved motif that regulates PZLS-1 abundance through a mechanism requiring DSH-1 and conserved, potentially phosphorylatable residues within the LNX3H domain. They reveal an additional level of PZLS-1 regulation downstream of DSH-1 and suggest that mechanisms controlling LNX3H-protein abundance may be evolutionarily conserved.

### PZLS-1 and DSH-1 regulate VAB-8 abundance and polarised localisation

In vertebrates, the atypical kinesin KIF26B has also been linked to Wnt-Ror-Dvl signalling. Wnt5a-Ror activation promotes KIF26B degradation, while KIF26B contributes to cell polarisation during directional migration^22,23,56^. This regulation parallels that of PDZRN3, although PDZRN3 and KIF26B can each respond to Wnt5a independently, indicating that the two branches are not strictly interdependent^23^. We therefore asked whether VAB-8, the *C. elegans* ortholog of KIF26B, contributes to Wnt-Ror-Dvl-dependent muscle polarity.

Complete deletion of *vab-8* did not disrupt TWK-28 polarity (Fig. 8A), indicating that VAB-8 is not required to establish this polarity pattern. However, VAB-8 was previously reported to display an asymmetric distribution in head body wall muscles^57^. We therefore generated a C-terminal endogenous tag common to all VAB-8 isoforms. VAB-8 was asymmetrically distributed throughout the body wall musculature, where it was restricted to the anterior tip of muscle cells (Fig. 8B and S6A). Its abundance was highest in head muscles and markedly lower in more posterior body wall muscles.

**Figure 8.**
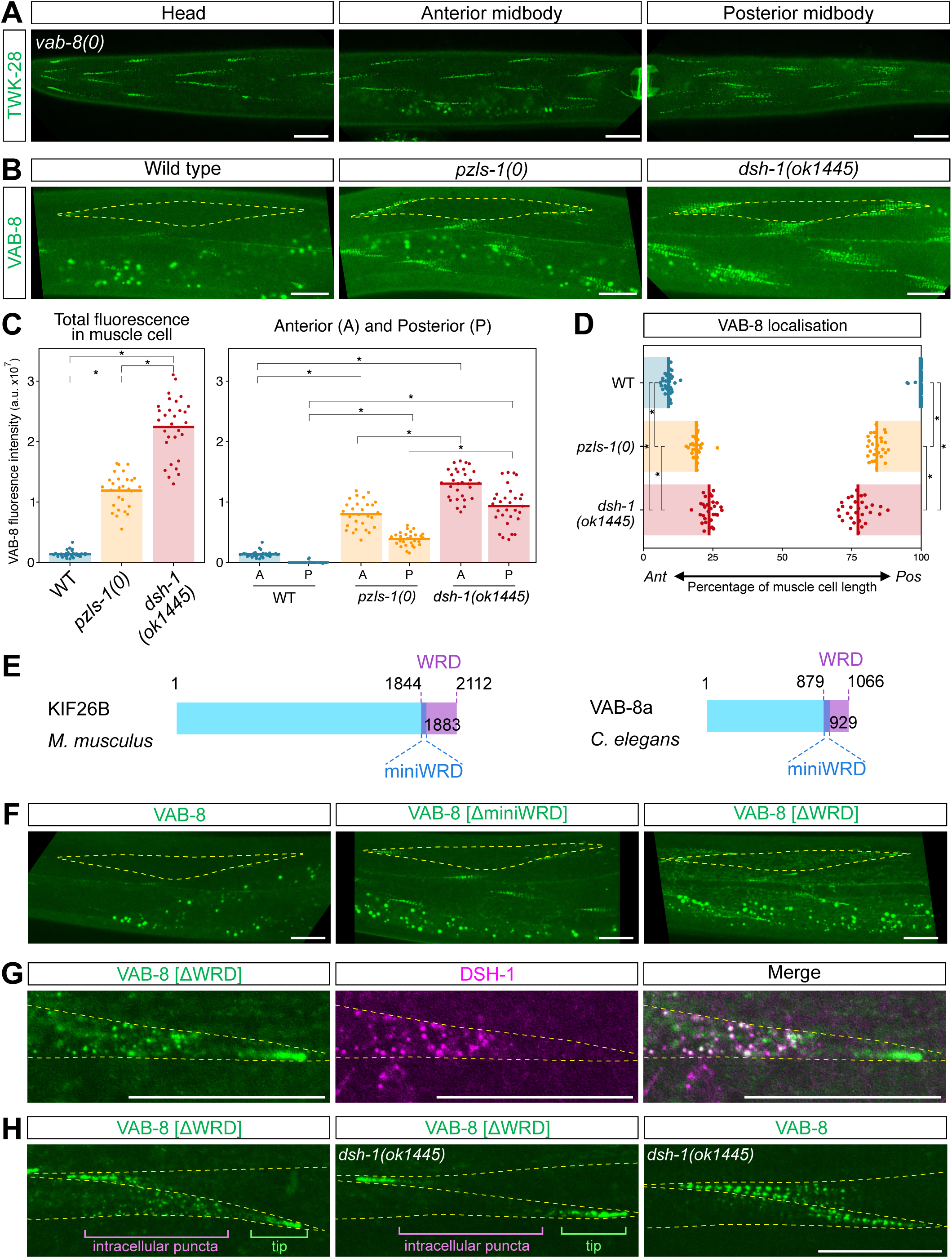
VAB-8/Kif26b abundance and localisation are regulated by DSH-1 and PZLS-1. **(A)** VAB-8 loss-of-function does not affect TWK-28 asymmetry. **(B)** VAB-8 is found at the anterior tip of body wall muscles. In *dsh-1* and *pzls-1* loss-of-function mutants, VAB-8 is present at both cell extremities and its abundance is increased. **(C-D)** Quantification of VAB-8-mNeonGreen fluorescence (*C*) and VAB-8 domain boundary positions (*D*). Coloured bars in *D* indicate the region containing VAB-8. Bars represent means; number of worms assayed for each condition: WT *n* = 31, *pzls-1(0) n* = 28, *dsh-1(0) n* = 30; *N* = 2 independent experiments; Kruskal-Wallis test followed by Dunn’s test with Holm’s correction * p<0.01. **(E)** Schematic representation of *M. musculus* KIF26B and *C. elegans* VAB-8a, indicating the WRD and miniWRD regions. Amino acid positions are shown in black. **(F)** Deletion of the miniWRD increases mNeonGreen-VAB-8 abundance without altering its localisation, whereas deletion of the entire WRD increases abundance and redistributes the truncated protein to both ends of muscle cells. **(G)** WRD-deleted VAB-8 protein forms intracellular puncta that colocalise with DSH-1 in the posterior end, but not the tip, of muscle cells. **(H)** Intracellular puncta formed by WRD-deleted VAB-8 are no longer detected in *dsh-1(0)* mutants, and WRD-deleted VAB-8 displays a pattern distinct from that of wild-type VAB-8 in a *dsh-1(0)* context. Muscle cell outlines are indicated with yellow dashed lines. Anterior to the left. Scale bars, 20 µm. Raw data and statistical analyses are presented in the Source Data file.

We next asked whether this polarised distribution depended on PZLS-1 and DSH-1. Loss of either protein caused VAB-8 to accumulate at both ends of muscle cells rather than exclusively at the anterior tip (Fig. 8B-D). VAB-8 fluorescence at cell extremities was also strongly increased in both mutant backgrounds, with a larger increase in *dsh-1* than in *pzls-1* mutants (Fig. 8C). Thus, PZLS-1 and DSH-1 regulate both the polarised localisation and abundance of VAB-8. The stronger effect observed upon loss of DSH-1 suggests either that additional factors act alongside PZLS-1 to control VAB-8, or that the defects of *pzls-1* mutants result indirectly from a partial disruption of DSH-1-dependent regulation of VAB-8.

In vertebrate cells, KIF26B contains a C-terminal region of approximately 270 amino acids, termed the Wnt5A-responsive degradation (WRD) domain, that is necessary and sufficient for Wnt5a-induced degradation^56^. A shorter sequence of approximately 40 amino acids within the WRD is required for this response; we refer to the homologous sequence in VAB-8 as the miniWRD (Fig. 8E and S6B). We next asked whether these conserved C-terminal regions similarly control VAB-8 abundance.

Deletion of the miniWRD from an endogenously tagged VAB-8 increased its fluorescence but did not alter its asymmetric localisation (Fig. 8F). This indicated that the miniWRD contributes to the regulation of VAB-8 abundance but is not required for its polarised distribution. Deletion of the entire WRD region also increased VAB-8 abundance, but in addition disrupted its localisation, with the truncated protein now accumulating at both ends of muscle cells (Fig. 8F). Importantly, this redistribution was not accompanied by a general loss of muscle polarity, as SGCB-1/β-Sarcoglycan remained asymmetrically localised in the same animals (Fig. S6C). Thus, the VAB-8 C-terminal region regulates both protein abundance and polarised localisation, with the miniWRD contributing specifically to abundance control.

WRD-deleted VAB-8 accumulated at both extremities of muscle cells. In addition, the truncated protein formed intracellular puncta near the posterior end of the cell that partially colocalised with DSH-1 (Fig. 8G). These intracellular puncta were no longer detected in *dsh-1* null mutants (Fig. 8H), indicating that their formation or maintenance depends on DSH-1. Importantly, terminal enrichment of WRD-deleted VAB-8 remained detectable in the absence of DSH-1, indicating that accumulation at cell extremities and formation of intracellular puncta are separable features of the mutant protein. The terminal enrichments formed by WRD-deleted VAB-8 were also shorter and denser than the broader VAB-8 accumulation observed in *dsh-1* or *pzls-1* mutants, and additional intracellular fluorescence was detected (Fig. 8F-H). Because deletion of the entire WRD could alter protein conformation or trafficking, the precise nature of these intracellular structures remains uncertain. Nevertheless, WRD-deleted VAB-8 retains the ability to accumulate at cell extremities while losing its normal restriction to the anterior tip, and its association with DSH-1-positive intracellular puncta is DSH-1-dependent.

The relationship between PDZRN3 and KIF26B remains incompletely understood in vertebrates. We therefore examined whether VAB-8 reciprocally influences PZLS-1 *in vivo* in *C. elegans*. Although loss of PZLS-1 strongly affected VAB-8 localisation and abundance, neither complete loss of VAB-8 nor deletion of the miniWRD or entire WRD detectably altered PZLS-1 localisation or abundance (Fig. S6D-F). Thus, VAB-8 is dispensable for PZLS-1 localisation and accumulation. Together with the effects of *pzls-1* loss on VAB-8, these findings support a unidirectional relationship in which VAB-8 functions downstream of PZLS-1 in this regulatory pathway.

Overall, these findings reveal a conserved relationship between Wnt-Ror-associated signalling and the control of KIF26-family proteins. In *C. elegans* muscles, PZLS-1 and DSH-1 regulate both VAB-8 abundance and polarised localisation, while conserved sequences within the VAB-8 C-terminus make separable contributions to these two properties.

## Discussion

In this work, we identify PZLS-1 as a new regulator of muscle polarity in *C. elegans*. PZLS-1 is the sole ortholog of the vertebrate LNX3H proteins PDZRN3, PDZRN4 and PDZD4/LNX5 and functions within a conserved non-canonical Wnt-Ror-Dvl signalling pathway. We show that PZLS-1 is itself asymmetrically localised *in vivo* and that its polarised distribution depends on EGL-20/Wnt, CAM-1/Ror and DSH-1/Dishevelled. Reciprocally, PZLS-1 is required for the polarised localisation of DSH-1, and the two proteins colocalise in a population of intracellular puncta, revealing a clear interdependence between these two components of the pathway. Through structure-function analyses, we identify conserved protein sequences that regulate PZLS-1 localisation, abundance and activity. We further place VAB-8, the *C. elegans* ortholog of the kinesin-like protein KIF26B, downstream of PZLS-1 in the context of muscle polarisation. Together, these findings support a model in which Wnt-Ror signalling establishes intracellular asymmetry through the coordinated spatial organisation and regulation of DSH-1 and PZLS-1, thereby contributing to the formation of highly reproducible polarised membrane compartments in *C. elegans* muscle cells.

In vertebrates, endothelial PDZRN3 is required for the oriented organisation of intermediate capillaries in the retina, and contributes to Wnt5a-induced directional cell migration *in vitro*^24^. Similarly, PDZRN3 overexpression disrupts the localisation of Cx43 and ZO-1 at cardiomyocyte intercalated disks^58^. Together with our findings in *C. elegans* muscles, these observations suggest that control of cellular polarity and polarised membrane organisation may represent an ancestral function of LNX3H-containing proteins.

Importantly, we show that PZLS-1 is itself asymmetrically distributed *in vivo*. To date, little is known about the subcellular localisation of LNX3H-containing proteins. Most studies have examined their localisation in cultured cells, including cancer cell lines^24,40,41,59,60^, or at the neuromuscular junction^27^ often following protein overexpression. To our knowledge, their localisation has not been investigated in a physiologically polarised cellular context. Notably, in our experiments, perturbations that altered PZLS-1’s asymmetric localisation were consistently associated with defects in muscle cell polarity, raising the possibility that spatial restriction of LNX3H proteins could be a key determinant of their function. Examining the endogenous localisation of PDZRN3 and PDZRN4 during polarised morphogenesis processes and in migrating cancer cells should therefore reveal whether asymmetric localisation is a conserved feature of this protein family.

In vertebrates, Wnt5a-Ror signalling promotes PDZRN3 degradation^23^. In contrast, our results argue that EGL-20/Wnt does not primarily regulate PZLS-1 by controlling its abundance. Two observations support this conclusion. First, muscle polarity can be restored across a range of PZLS-1 expression levels, indicating that its function is not tightly coupled to absolute protein abundance. Second, loss of PZLS-1 phenocopies loss of EGL-20/Wnt, with disruption of either component leading to loss of polarity. This is difficult to reconcile with a model in which EGL-20/Wnt establishes polarity mainly by promoting PZLS-1 degradation. Instead, our data support a model in which EGL-20-CAM-1 signalling primarily controls the spatial organisation of PZLS-1. Thus, Wnt-Ror signalling may regulate LNX3H-family proteins through distinct, context-dependent mechanisms, including control of both protein abundance and subcellular localisation.

This possibility is particularly interesting in light of observations in vertebrate cells. *In vitro,*HUVEC cells exposed to a Wnt5a gradient undergo directional migration in a PDZRN3-dependent manner^24^. In this context, an asymmetric Wnt5a signal could locally reorganise, stabilise or destabilise PDZRN3 to generate intracellular asymmetry. On the other hand, the requirement for both Wnt5a and PDZRN3 for proper polarised migration is difficult to reconcile with a model in which Wnt5a acts solely by promoting PDZRN3 degradation.

Nevertheless, our data also indicate that PZLS-1 abundance is regulated. Deletion of the previously uncharacterised abundance regulatory domain (ARD) caused a marked increase in fluorescence of the endogenous PZLS-1 translational reporter, and this effect depended on DSH-1. Notably, this short conserved motif is also present in vertebrate LNX3H proteins, raising the possibility that it contributes to the regulation of their abundance, potentially in the context of Wnt5a-dependent control of PDZRN proteins. Thus, non-canonical Wnt signalling may regulate LNX3H proteins at multiple levels, coordinating both their spatial organisation and abundance.

PDZRN3 is phosphorylated on conserved residues upon Wnt5a signalling, and this phosphorylation is crucial for its subsequent degradation^23^. Here, we show that these conserved serine/threonine residues are also required for PZLS-1 function. Yet, it is again unlikely that they are required to modulate PZLS-1 levels. Instead, they may regulate protein conformation or interfere with protein-protein interactions. Consistently, these residues are also conserved in PDZD4, which is not degraded by Wnt5a signalling^23^. Our results thus suggest that these S/T residues may support both degradation and non-proteolytic functions of LNX3H-containing proteins.

PDZRN3 and PDZRN4 have been studied predominantly in the context of their RING finger-dependent E3 ubiquitin ligase activity. However, as for PZLS-1, many LNX3H-family proteins and isoforms lack a RING finger domain, both in vertebrates and across metazoan lineages^26,61^. The functions of these RING-less proteins remain largely unexplored. Notably, a recent study reported that mouse skeletal muscle predominantly expresses a PDZRN4 isoform lacking the RING finger domain and that this isoform is required for neuromuscular junction maintenance^27^. These observations suggest that important functions of LNX3H proteins are independent of their intrinsic ubiquitin ligase activity. We therefore propose that PZLS-1 and *C. elegans* muscle polarity provide a useful *in vivo* model for dissecting the conserved, RING-independent functions of LNX3H-domain proteins.

A major next step will be to define the molecular partners and assemblies through which PZLS-1 functions. In vertebrates, PDZRN3 has been reported to interact with Dvl3 and to promote Dvl internalisation in response to Wnt signalling through its ubiquitin ligase activity^24^. Here, we find that endogenously tagged PZLS-1 and DSH-1 colocalise within muscle cells and are reciprocally required for their polarised localisation. These observations point to a close functional association between the two proteins in the context of Wnt signalling and, importantly, show that such an association can occur in the absence of a RING finger domain and its associated ubiquitin ligase activity.

The nature of the intracellular puncta containing both PZLS-1 and DSH-1 remains unclear. They did not correspond to the canonical endosomal compartments examined here. Dishevelled proteins have long been known to form intracellular puncta, initially in studies using overexpressed reporters^49,50,62^ and, more recently, at endogenous expression levels^63^. Dvl2 can also undergo liquid-liquid phase separation, a property implicated in canonical Wnt signalling^63^. One possibility is therefore that PZLS-1 associates with DSH-1 in biomolecular condensates. Testing this hypothesis will require determining the dynamics and biophysical properties of these puncta, as well as assessing the contribution of intrinsically disordered regions to their formation.

The partial colocalisation of EGL-20/Wnt with some of these puncta adds another layer of complexity. Wnt ligands can be internalised by responding cells^7,64^, and receptor-bound Wnt signalling complexes have been proposed to undergo internalisation^15^, which might instead suggest a membrane-associated compartment. Conversely, the Wnt co-receptor LRP6 has also been detected in endogenous Dvl2-containing condensates^65^, indicating that the presence of membrane-associated signalling components does not necessarily exclude a condensate-like organisation. Resolving the molecular composition, membrane association and dynamics of the PZLS-1-DSH-1 puncta will therefore be important for understanding how these structures are formed, and how they may contribute to Wnt-Ror signalling in muscle cells.

This relationship may extend to VAB-8/KIF26. A C-terminally truncated form of VAB-8 colocalised with DSH-1 in intracellular puncta, reminiscent of observations in vertebrate cells in which KIF26B colocalises with Dvl3 following overexpression^25^. KIF26B, the vertebrate ortholog of VAB-8, is itself regulated by non-canonical Wnt signalling: Wnt5a-Ror-Dvl signalling promotes its proteasome-dependent degradation^22,56^. Here, we show that DSH-1 and PZLS-1 regulate both the abundance and polarised localisation of VAB-8. By contrast, loss or truncation of VAB-8 did not detectably affect PZLS-1 abundance or localisation, supporting a directional relationship in which VAB-8 functions downstream of PZLS-1. This provides an *in vivo* framework for understanding the potential crosstalk between PDZRN- and KIF26-family proteins downstream of Wnt-Ror signalling^23^. Moreover, deletion of the VAB-8 region homologous to the KIF26B Wnt5A-responsive degradation domain increased VAB-8 abundance, suggesting that conserved C-terminal determinants contribute to the regulation of both proteins. The broader VAB-8 C-terminal region was also required for its restriction to the anterior tip of muscle cells, providing an entry point for dissecting how KIF26-family proteins become spatially organised within polarised cells.

Despite this conserved regulation, VAB-8 itself appears dispensable for the establishment of muscle polarity. This contrasts with vertebrate KIF26B, which contributes to directional cell migration in response to Wnt signalling^25^. One possibility is that VAB-8 acts redundantly with other downstream effectors. Consistent with this idea, *vab-8* loss-of-function causes only partial and variable neuronal migration defects and has been proposed to act redundantly with other factors in these cells^66^. An alternative possibility is that the relevant output of Wnt-Ror signalling is not simply activation of VAB-8, but rather the spatial restriction of its abundance and localisation. Indeed, when DSH-1 or PZLS-1 function is disrupted, VAB-8 accumulates at both muscle-cell extremities and its abundance increases. Distinguishing between these possibilities will require perturbations that selectively alter VAB-8 localisation while preserving overall protein function. Deletion of the entire WRD removes a substantial fraction of VAB-8 and is therefore unlikely to affect localisation alone: the truncated protein displays a distribution distinct from that of full-length VAB-8 in polarity-defective backgrounds and shows additional intracellular accumulation, potentially reflecting altered trafficking or protein organisation. A systematic structure-function analysis should therefore help define smaller determinants specifically required for VAB-8 polarised localisation, while minimising broader effects on protein structure and function.

A major unresolved question is how PZLS-1 activity is translated into muscle cell polarisation. The downstream effectors of PDZRN3 remain poorly defined, although PDZRN3 has been linked in some contexts to PKCζ and c-Jun signalling^24,34,58^. More broadly, several studies have implicated JNK and c-Jun downstream of Wnt5a-Ror signalling^13,67–69^, although this connection appears context-dependent, as c-Jun phosphorylation was unaffected in Wnt5a^−/−^ or Ror1^−/−^;Ror2^−/−^ fibroblasts^14^. Whether these pathways contribute to muscle polarisation in *C. elegans* remains unknown. At the cellular level, PZLS-1 may ultimately influence polarity through cytoskeletal reorganisation and/or directed intracellular transport. Consistent with a potential role in cytoskeletal regulation, PDZRN3 is required for F-actin stress-fibre formation following serum deprivation *in vitro*^34^, whereas PDZRN3 overexpression in mouse myocardium increases the G-actin/F-actin ratio^58^.

In this context, identifying the molecular partners of PZLS-1 will be particularly informative. Defining the interactome of PZLS-1 and its downstream effectors should ultimately help bridge the gap between Wnt-Ror-Dvl signalling and the reproducible asymmetric organisation of muscle cells. Furthermore, as discussed above, multiple LNX3H-family proteins and isoforms lack a RING finger, indicating that the functions of this protein family are unlikely to be restricted to E3 ubiquitin ligase activity. Identifying PZLS-1 interactors could uncover molecular functions encoded by the evolutionarily conserved LNX3H core, independently of the RING finger domain.

Together, our results suggest that PDZRN proteins may have conserved roles in cellular polarisation, and *C. elegans* PZLS-1 provides a genetically tractable model to study *in vivo* how Wnt-Ror can signal via LNX3H proteins to organise cell polarity.

## Methods

### *C. elegans* methods and genetics

*C. elegans* cultivation, genetic crosses and manipulation were carried out according to standard protocols^70^. Unless otherwise specified, *C. elegans* strains were maintained at 20°C, and were grown on nematode growth medium (NGM) agar plates seeded with *Escherichia coli* OP50 as a food source. All *C. elegans* strains were originally derived from the wild-type Bristol N2 strain. Strains and alleles used for this study are listed in Supplementary Tables 1 and 2, respectively.

### Mutagenesis

Mutagenesis was performed on the strain JIP5526. Worms were collected and washed with M9 buffer (3 g of KH_2_PO_4_, 6 g of Na_2_HPO_4_, 5 g of NaCl, and 0.25 g of MgSO_4_·7H_2_O, distilled water up to 1 liter), mutagenesis was performed by adding 20 μl of EMS (Sigma-Aldrich, M-0880) to 4 mL of M9 containing the pellet of worms, for 4 h at 20°C with rotation. Worms were then washed ten times in M9 and put back on NGM plates with OP50 (4 L4 per plate). They were then grown at different temperatures to enable differential growth timings. F1 offspring were cloned, and when F2 worms reached the adult stage they were mounted on slides with 2% (w/v in water) dry agarose pads, immersed in 2 % polystyrene beads (Polybeads Microsphere 0.10 µm, Polysciences, reference 00876) in M9 buffer. Fluorescence was observed live under an AZ100 macroscope (Nikon) equipped with a Flash 4.0 CMOS camera (Hamamatsu Photonics). Mutant worms with a loss of polarity were isolated and bred to homozygosity. Genomic DNA was extracted from homozygous strains with the DNeasy blood and tissue kit (Qiagen, reference 69506) and sent for whole-genome sequencing (Eurofins). EMS-induced mutations were identified using in-house built scripts.

### CRISPR/*Cas9* genome engineering

All knock-in alleles were generated according to Ghanta and Mello^71^. crRNAs were designed with Benchling software and ordered from Integrated DNA Technologies (IDT). Injection mixes were prepared with 2.8 μL of crRNA (34 μM), 5 μL of tracrRNA (18 μM, IDT #1073189) and 1 μL of Cas9 nuclease (5 µg/µl, Sigma) incubated for 15 min at 37 °C. For the generation of deletions, two crRNAs (1.4 μL each) were used, each crRNA targeting an end of the target sequence. After heating, 800 ng of the pRF4 plasmid was added, as well as the repair template: for point mutations or deletions, the repair template was 2.2 μl of a single-strand oligonucleotide (1 μg/μl), and for insertions larger than 130 bp, either 500 ng of melted double-strand DNA ^71^ or 660 ng of single-strand DNA prepared as described by Eroglu *et al*.^72^. IDT nuclease-free water was added to the mix up to 20 μl. Adult hermaphrodites were injected with this mix, and in the next generation plates containing worms with a roller phenotype were selected. Roller or non-roller F1 were isolated and tested by polymerase chain reaction (PCR) for the presence of the desired modification. All edited sequences were verified by Sanger sequencing.

### Molecular biology

Single-strand oligonucleotides and crRNAs used in this study are described in Supplementary Tables 3 and 4, respectively. The *myo-3* promoter was used for tissue-specific expression in body wall muscles. Plasmid sequences are available upon request, and all constructs were verified by Sanger sequencing. Plasmids constructed for this study are described in Supplementary Table 5.

### Generation of single-copy insertion alleles

Single-copy insertions were generated using the miniMos approach^73^. Briefly, adult *C. elegans* hermaphrodites were injected with a mix composed of 15 ng/μL of miniMos plasmid, 50 ng/μL of pCFJ601 plasmid (*Mos1* transposase), 10 ng/μL of pMA122 plasmid (negative selection marker *Phsp16.2::peel-1*), and 2.5 ng/μL of pCFJ90 plasmid (*Pmyo-2::mCherry*). Neomycin (G418) was added to plates 24-48 h after injection at 1 μg/μL final concentration. Candidate plates were heat-shocked for 2 h at 34°C. Worms with an insertion were isolated and bred to homozygosity, and the absence of extrachromosomal arrays containing the pCFJ90 plasmid was verified by confirming the absence of red fluorescence in pharynx muscles.

### Confocal imaging

For live *in vivo* imaging, hermaphrodites around the young adult stage were mounted on 2% (w/v in water) dry agarose pads in 2 % polystyrene beads diluted in M9 buffer. Confocal images were acquired on Andor Revolution Imaging System (Oxford Instruments). The spinning disk imaging system was composed of an inverted microscope (Olympus IX83) equipped with 40x/NA 1.3 and 60x/NA 1.4 oil immersion objectives, a Yokogawa CSU-X confocal scanning unit and an EMCCD camera (iXon Ultra 888 Andor). Acquisition settings were identical across genotypes for quantitative or qualitative analysis. Images were obtained with Metamorph software (version 7.10.5.476) as a stack of optical sections (0.2 μm apart). Images shown are maximum-intensity projections along the Z-axis.

### Fluorescence quantification

Image analysis was performed with Fiji (ImageJ2 version 2.3.0/1.53q). Each data point represents one muscle cell, and only a single muscle cell was quantified per worm. Images for quantifications were acquired in the anterior midbody of the worms. ROIs were drawn blind to the genotypes, when possible (in some cases the clear difference between wild-type or mutant patterns made it impossible to be blinded to genotype).

*Quantification of TWK-28, SLO-1 and VAB-8 localisation domains and PZLS-1 posterior enrichment length –* A 40x objective was used, allowing to see the entire length of the cells. ROIs were drawn using TWK-12::wrmScarlet or PAT-2::BFP to outline muscle membrane. The lengths of anterior or posterior domains were normalised over the length of the associated cell. *Quantification of TWK-28 and VAB-8 fluorescence intensities –* A 40x objective was used, allowing to see the entire length of the cells. 5 confocal planes centred on the best focus plane were summed, and ROIs were drawn using TWK-12::wrmScarlet or PAT-2::BFP to outline the muscle membrane. Fluorescence intensities were corrected by subtracting background fluorescence measured in the middle of the cell, where TWK-28 or VAB-8 were undetectable. For TWK-28, the corrected fluorescence intensities values were normalised over the mean total fluorescence in wild type from data of the same day, due to variations across days.

*Quantification of PZLS-1 fluorescence intensities in the posterior enrichment –* A 60x objective was used. 5 confocal planes centred on the best focus plane of the surface posterior enrichment of PZLS-1 were summed, and ROIs were drawn using TWK-12::wrmScarlet to outline the muscle membrane. For the quantification of ARD mutants, only a single focal plane was used. The background fluorescence was estimated outside muscle cells, within the worm, and was used to correct fluorescence intensity values.

### Statistics and reproducibility

Graphical representations and statistical analyses were generated with R (R 4.2.1) used with RStudio (version 2023.09.1+494). Statistical analyses are reported in the Source Data file. In mutant animals, polarity was assessed in >30 animals for each genotype.

### Protein sequences and alignments

Protein sequences used in alignments for this study were retrieved from WormBase, UniProt or GenBank. All accession identifiers are available in Supplementary Table 6. Alignments were made with Jalview, using mafft alignments with default parameters, except the gap opening penalty that was set to 1 for PZLS-1/PDZRN alignments.

Sequences used for schematics of human isoforms shown in Figure 1B are as follows PDZRN3: NP_055824.1, NP_001290068.1, NP_001290070.1, NP_001290071.1. PDZRN4: NP_001158067.1, NP_037509.3. PDZD4: NP_001290441.1, NP_115901.2, NP_001290442.1, NP_001290443.1, NP_001290444.1, NP_001290445.1.

### Structure prediction and analysis

Protein complex structure predictions with Alpha-Fold-Multimer was performed with the ColabFold implementation version 1.5.2 running Alphafold release 2.3.1^74–76^. For the prediction settings, we used the default settings including the use of template information and relaxation of the predicted structures using amber force elds. iPTM value for the model in Figure S5 was 0.265. Analysis of the structures and models display was performed using UCSF ChimeraX version 1.7.1 (2024-01-23).

## Data availability

All strains and reagents are available upon request to T.B.

## Acknowledgements

We thank members of the Boulin and Bessereau labs, Julien Falk, Marie Gendrel and Martin Harterink for providing feedback, and Driss Laabid for technical assistance. This work was funded by a grant to T.B. from the French National Research Agency (ANR) (AAPG 2023, DYSCO). E.C. was supported by a PhD fellowship from the French Ministry of Research (MESRI) and Fondation pour la recherche médicale (FRM). This work was performed within the framework of the LABEX CORTEX (ANR-11-LABX-0042) of Université Claude Bernard Lyon 1, within the programme “Investissements d’Avenir” (decision no. 2019-ANR-LABX-02) operated by the French National Research Agency (ANR). Additional funding was provided by the Université Claude Bernard Lyon 1, CNRS (Centre National de la Recherche Scientifique) and Inserm (Institut national de la santé et de la recherche médicale). We thank the Caenorhabditis Genetics Center (CGC), funded by NIH Office of Research Infrastructure Programs (P40 OD010440), for providing strains. We thank the SFR Santé Lyon-Est core facility (UAR3453 CNRS, US7 Inserm, UCBL), specifically the CIQLE platform, a member of LyMIC and the national infrastructure France-BioImaging, funded by the French National Research Agency under grant ANR-24-INBS-0005 FBI BIOGEN, for support and access to equipment and Camilla Luccardini for technical assistance. Some strains were generated by SEGiCel (SFR Santé Lyon Est CNRS UAR 3453, Lyon, France) with the support of CNRS and IBiSA, within the framework of Celphedia.

## Author contributions

Conceptualisation: E.C. and T.B., Methodology: E.C., S.D., T.B., Investigation: E.C and S.D., Writing—original draft: E.C Writing—review and editing: E.C. and T.B., Funding acquisition: T.B, Supervision: T.B.

---

**Supplementary Figure 1. PZLS-1/PDZRN regulates TWK-28 and SGCB-1 asymmetry in muscle and bears domains conserved in LNX3H proteins**

**(A)** Mutations in *pzls-1* disrupt the polarised localisation of SGCB-1/β-Sarcoglycan.

**(B)** Asymmetrical distribution of SGCB-1 in a wild-type context, and symmetrical localisation in a *pzls-1* loss-of-function, in the posterior midbody.

**(C)** Quantification of TWK-28 domain boundary positions in wild type, *dsh-1* and *pzls-1* loss-of-function mutants. Coloured bars indicate the region containing TWK-28. Bars represent data means; number of animals assayed: *n =* 77, 72, 56 (for respective rows); *N* = 3 independent experiments; Kruskal-Wallis followed by Dunn’s test with Holm’s correction. * p < 0.01.

**(D)** Quantification of TWK-28 abundance in anterior (A) or posterior domains (P) of individual muscle cells. Normalised fluorescence intensity relative to mean fluorescence of WT. Bars represent data means; number of animals assayed: WT *n =* 77, *pzls-1(0) n = 72*, *dsh-1(0) n =* 56; *N* = 3 independent experiments; Kruskal-Wallis followed by Dunn’s test with Holm’s correction. * p < 0.01. Muscle cell outlines are indicated with yellow dashed lines. Anterior to the left. Scale bars, 20 µm. Raw data and statistical analyses are presented in the Source Data file.

**(E)** Sequence alignment of LNX3H-containing proteins with the representation of conserved domains and predicted coiled-coil regions. Accession numbers are provided in Supplementary Table 1. Domain annotations are based on human PDZRN3 unless otherwise noted. The ring finger domain annotation is based on UniProt description of human PDZRN3 and PDZRN4. The TRAF type domain specific to PDZRN3 has not been annotated. PDZ domain annotations are based on UniProt description of the human proteins, and were slightly extended to encompass the entire PDZ domain based on AlphaFold structure predictions. The LNX3H domain is based on the definition by Flynn *et al*. 2011. The β-sheet annotation is based on the prediction by Flynn *et al.* 2011, and structure prediction by AlphaFold. Three residues in the LNX3H domain that are phosphorylated in Pdzrn3^77^ are indicated by red asterisks. Coiled-coil regions predicted by CoCoNat^78^ in human PDZRN3 are annotated in grey. Coiled-coil domains are also predicted for human PDZRN4 and PDZD4, and *C. elegans* PZLS-1 in these regions, with slight differences in the border of the domains. Prediction of the coiled-coil domain present in PZLS-1 was confirmed by deepcoil2^79^ prediction. The ARD domain annotation is based on sequence conservation and experimental data presented here.

**Supplementary Figure 2. *pzls-1* expression pattern.**

**(A)** *pzls-1* transcription levels in the 25 highest-expressing cell types. Based on adult hermaphrodite single-cell RNAseq data, all cells unfiltered (the CeNGEN Consortium*, in preparation*).

**(B)** *pzls-1* transcriptional reporter knock-in reporter line (*bab688*). Body wall muscles (red asterisks), vulval muscles (orange arrowhead), motor neurons (white arrowheads), and anal depressor muscles (pink arrowhead). *Left panel*, ventral view. *Right panel*, lateral view of tail region, ventral side on top.

**(C)** Single-copy muscle-specific transgenic expression lines with different mNeonGreen-PZLS-1 levels rescue SGCB-1 localisation in *pzls-1(0)*. Anterior to the left. Scale bars, 20 µm.

**Supplementary Figure 3. Intracellular localization of DSH-1 relative to EGL-20/Wnt.**

**(A)** Partial colocalization of EGL-20/Wnt with DSH-1 in anterior midbody muscle cells. *First row*, surface of muscle cells. *Second row*, interior of muscle cells. *Third row*, magnified views of the posterior end of a muscle cell. Anterior to the left. Scale bars, 20 µm.

**Supplementary Figure 4. Deletion of the DSH-1 PDZ domain disrupts TWK-28 polarity.**

**(A)** Deletion of the DSH-1 PDZ domain disrupts TWK-28 polarity.

**Supplementary Figure 5. Mutations in a predicted β-sheet within the LNX3H domain do not prevent PZLS-1 asymmetrical distribution in head muscles.**

**(A)** AlphaFold-Multimer prediction of a heteroduplex containing PZLS-1 (in orange, the PDZ binding motif is in magenta) and DSH-1e (in grey).

**(B)** PZLS-1 W286G and W286E variants retain an asymmetrical distribution in head muscle cells.

**(C)** *dsh-*1 loss-of-function or deletion of the predicted β-sheet impair the posterior enrichment of PZLS-1 in head muscles. Anterior to the left. Scale bars, 20 µm.

**Supplementary Figure 6. PZLS-1 localisation is independent of VAB-8**

**(A)** VAB-8 is asymmetrically distributed in body wall muscles and has a stronger expression in head muscles. Stitched mosaic of fluorescence images spanning the entire animal.

**(B)** Alignment of mouse KIF26B and *C. elegans* VAB-8a. miniWRD, blue. WRD, blue and purple.

**(C)** Deletion of the VAB-8 miniWRD or WRD does not disrupt the asymmetric localisation of SGCB-1/β-Sarcoglycan. Muscle cell outlines are indicated with yellow dashed lines.

**(D)** PZLS-1 localisation is unaffected by the complete deletion of *vab-8*, by deletion of VAB-8 WRD domain, or by deletion of the miniWRD domain. Muscle membranes are labelled by TWK-12-wrmScarlet (in magenta).

**(E)** Quantification of PZLS-1 posterior enrichment in a *vab-8* null background, or mutants lacking VAB-8 WRD or miniWRD domains. Coloured bars indicate the region containing PZLS-1; bars represent means; number of animals assayed in each condition: WT *n* = 32, *vab-8(0) n* = 31, *vab-8[ΔminiWRD] n* = 32, *vab-8[ΔWRD] n* = 38; *N* = 2 independent experiments; Kruskal-Wallis test, n.s., not significant.

**(F)** Quantification of PZLS-1 fluorescence in a *vab-8* null background, or mutants lacking VAB-8 WRD or miniWRD domains. Bars represent means; number of animals assayed in each condition: WT *n* = 30, *vab-8(0) n* = 29, *vab-8[ΔminiWRD] n* = 32, *vab-8[ΔWRD] n* = 27; *N* = 2 independent experiments; ANOVA, n.s., not significant.

Anterior to the left. Scale bars, 20 µm. Raw data and statistical analyses are presented in the Source Data file.

