## Supplementary Figures for "PZLS-1/PDZRN controls muscle polarity downstream of Wnt-Ror-Dvl signalling in *C. elegans* muscles"

#### Supplementary Figure 1 - 1/4

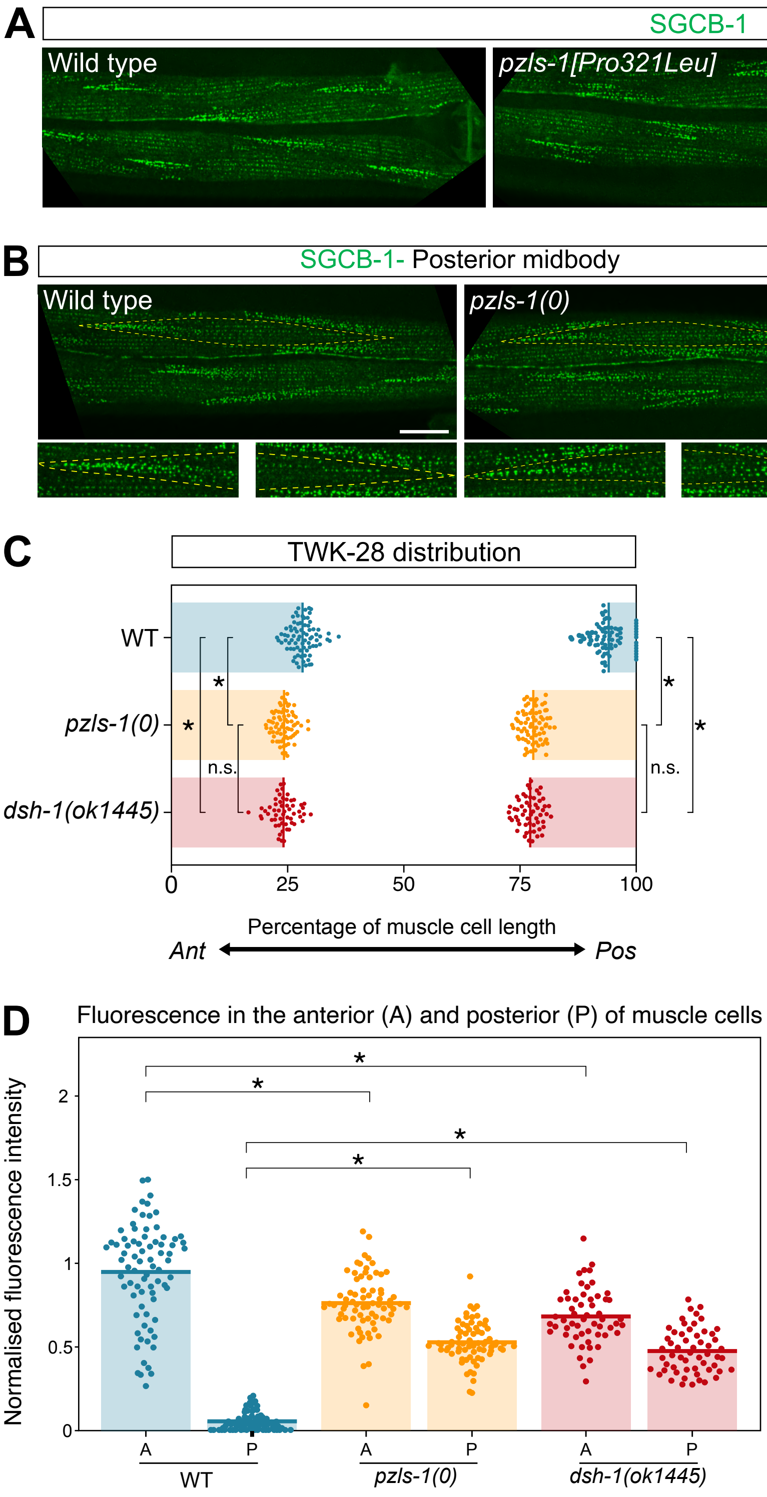

**Supplementary Figure 1 - 2/4**

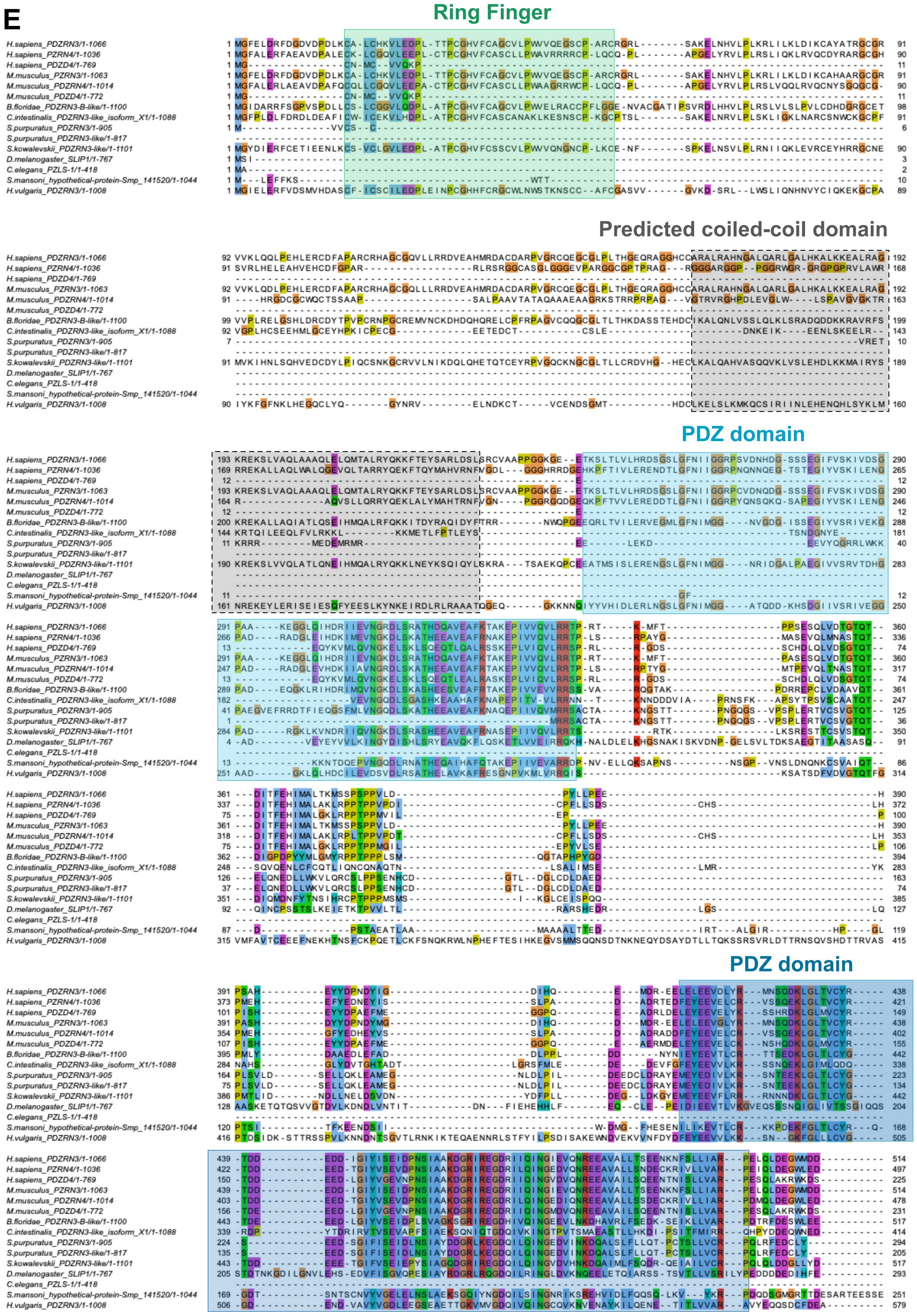

### Supplementary Figure 1 - 3/4

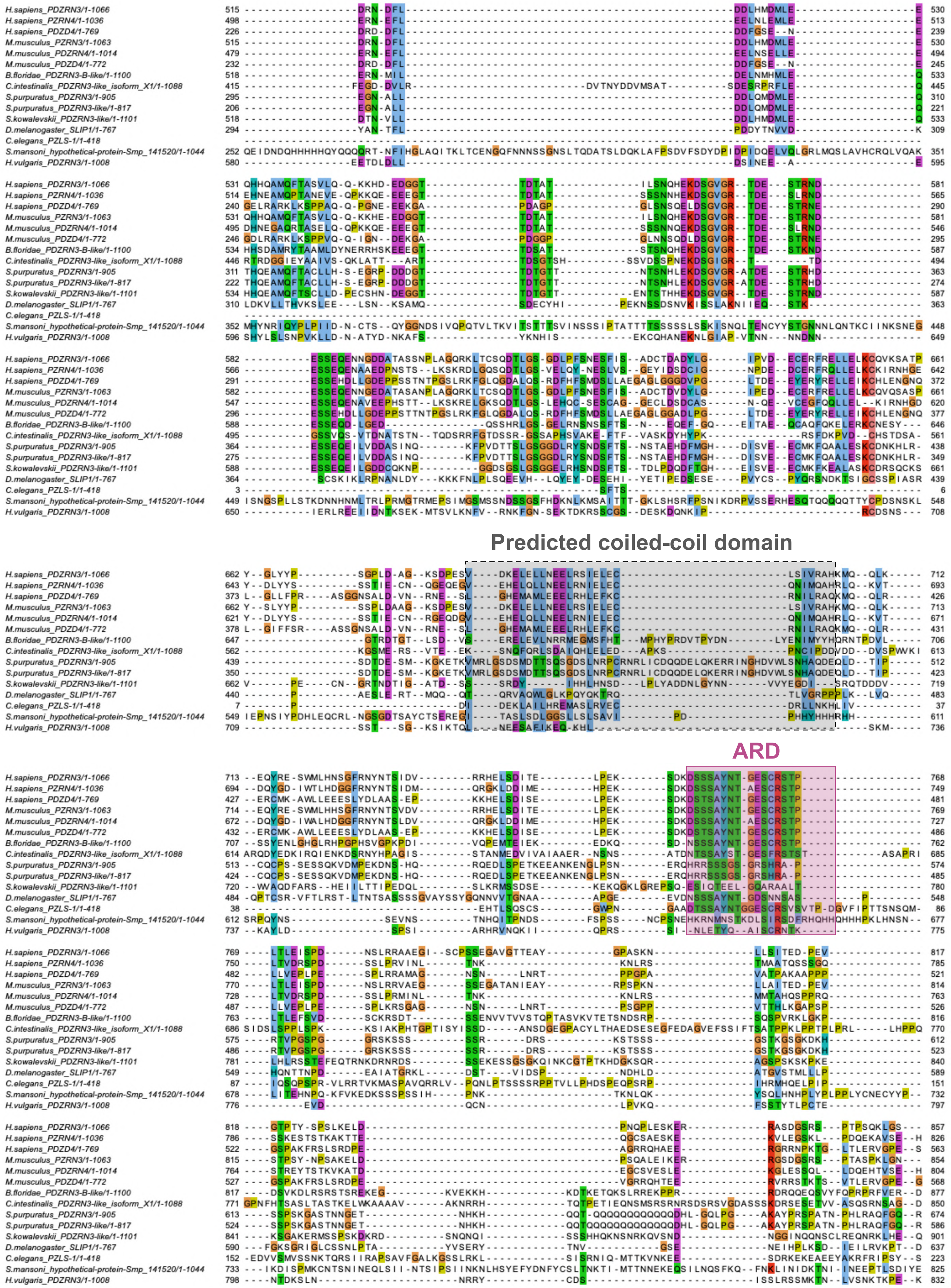

### Supplementary Figure 1 - 4/4

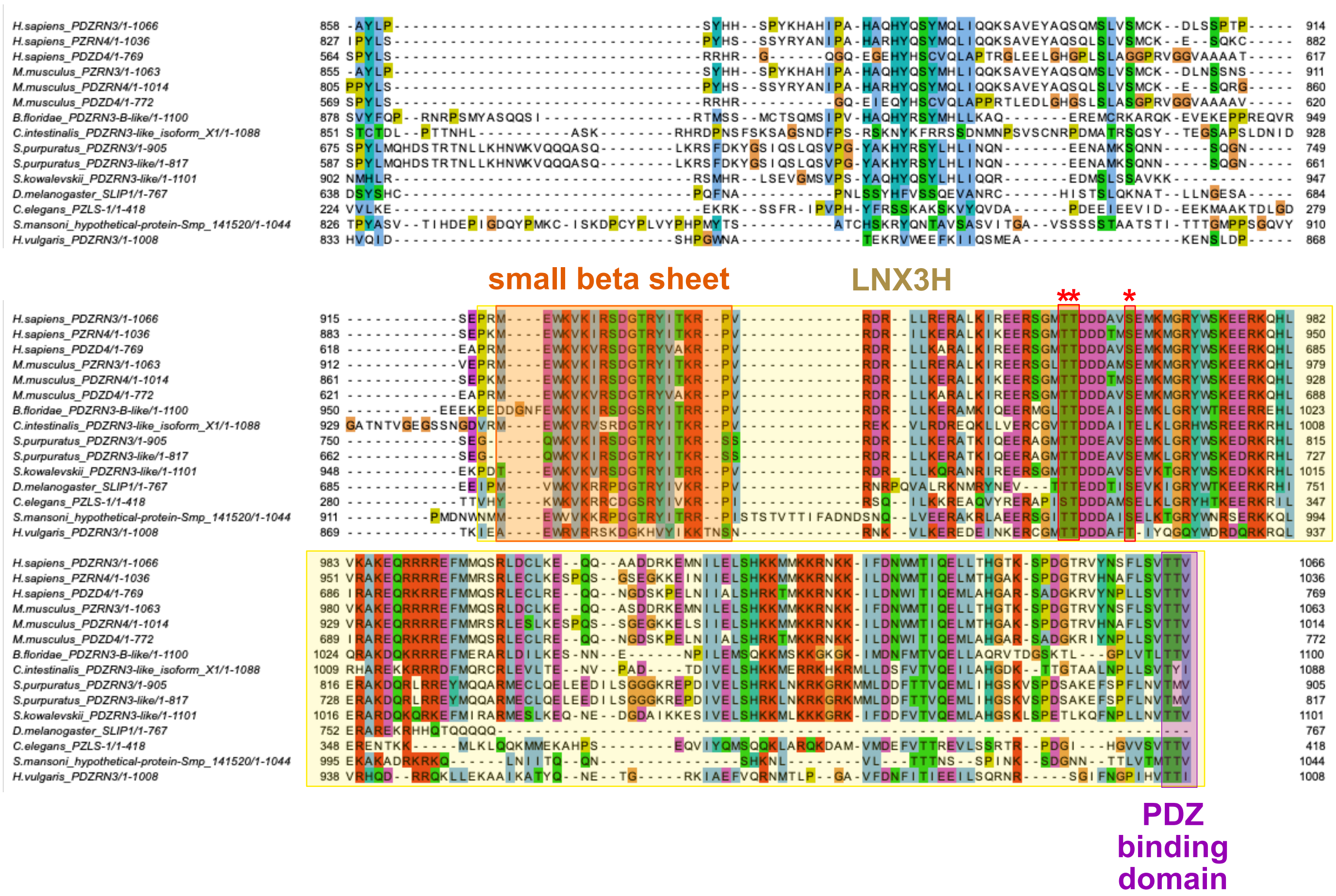

#### Supplementary Figure 2

**A**

| Cell type | Expression level |
|---|---|
| RIF | 262.701 |
| DA1 | 183.625 |
| DA | 170.211 |
| SAA | 121.703 |
| AS | 118.162 |
| Body_wall_muscle | 114.104 |
| PDB | 91.003 |
| DVB | 82.398 |
| Vulval_muscle_vm2 | 74.865 |
| DVC | 69.647 |
| DVA | 65.671 |
| AS11 | 65.24 |
| VA | 64.171 |
| PHC | 55.525 |
| Vulval_muscle_vm1 | 52.436 |
| Hypodermis_tail | 49.295 |
| VC_4_5 | 46.735 |
| AVF | 45.317 |
| SAB | 44.98 |
| PVP | 32.825 |
| VA1 | 31.838 |
| Anal_muscle | 30.923 |
| ASH | 27.796 |
| PHso_2? | 25.046 |
| AVG | 24.982 |

**B**

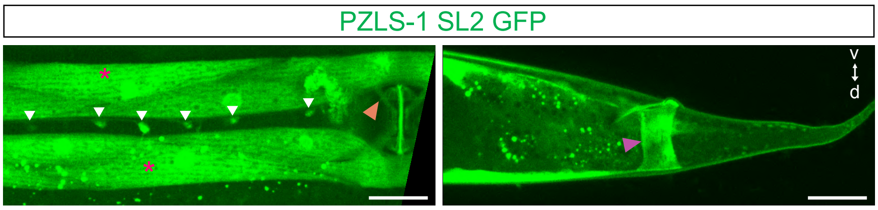


**C**

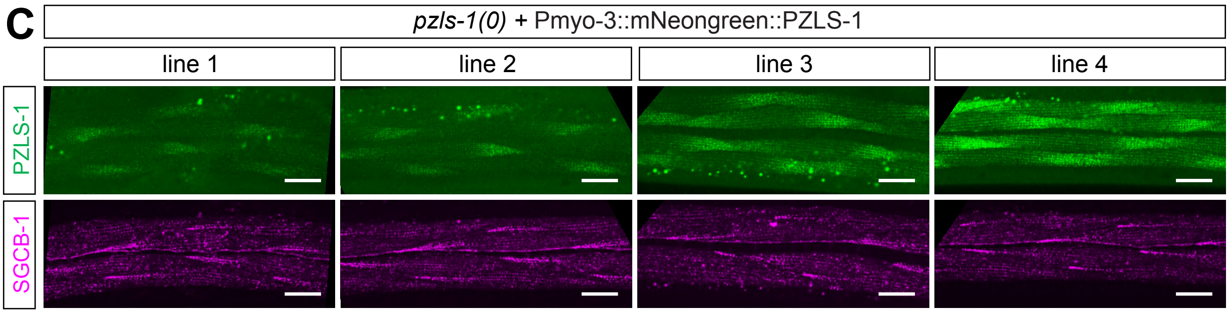

**Supplementary Figure 3**

#### Anterior to the vulva

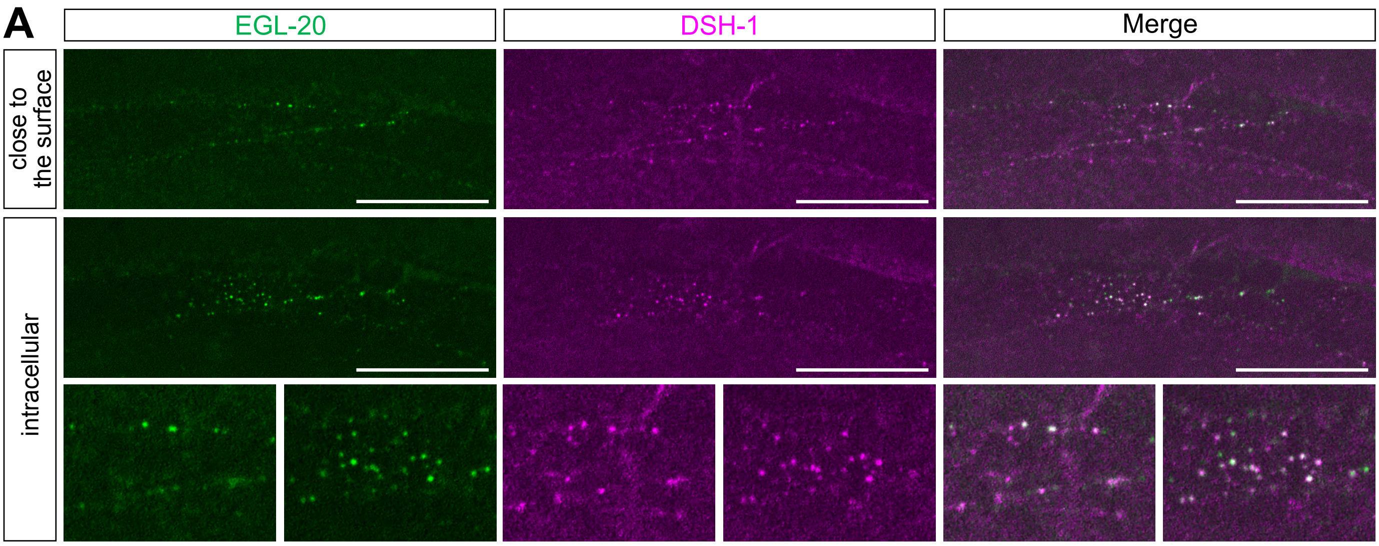

#### Supplementary Figure 4

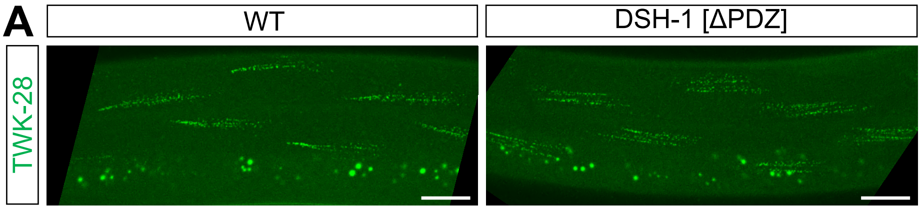

#### Supplementary Figure 5

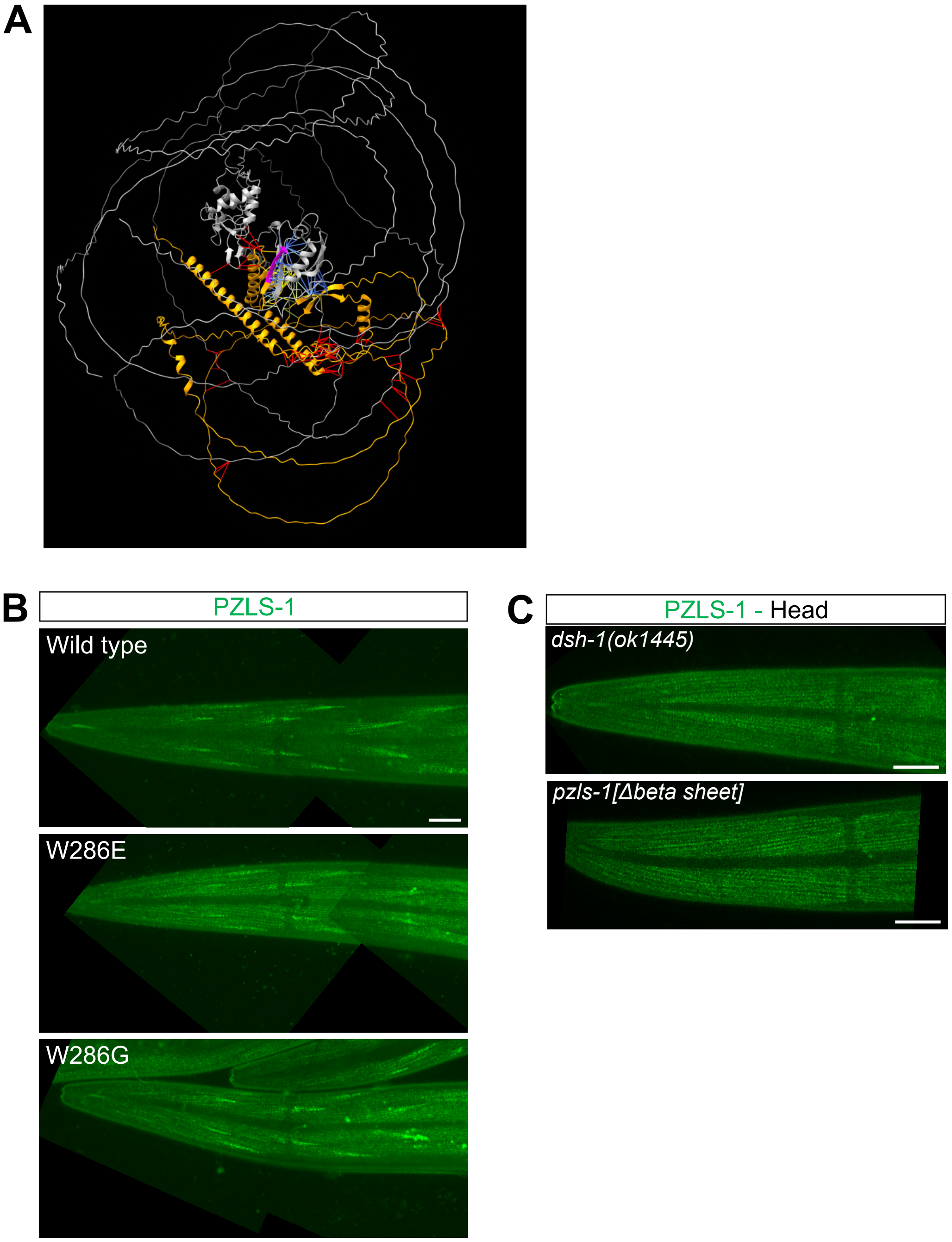

#### Supplementary Figure 6

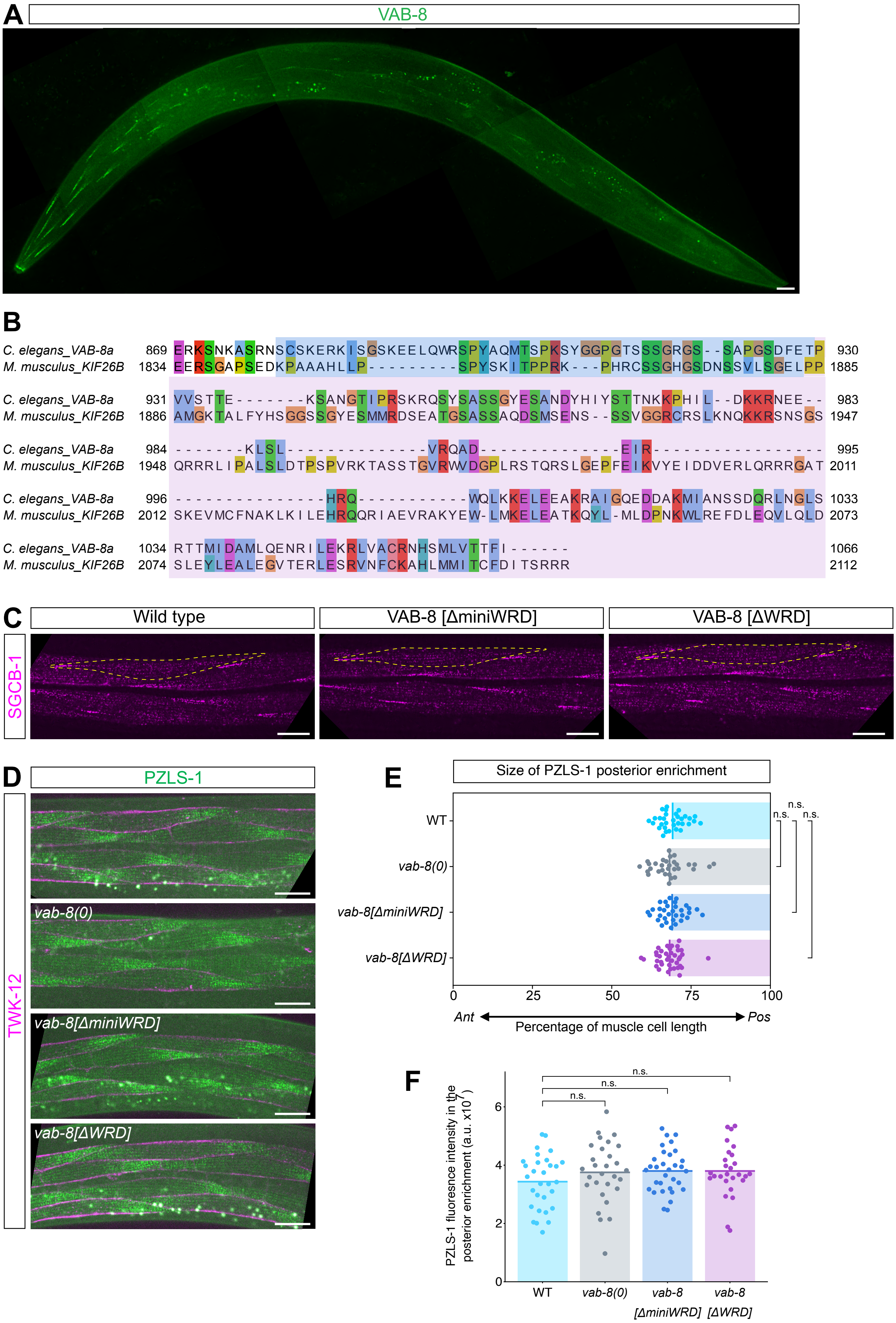
